# Metabolic Immunosuppression Mediated by Proliferative CD168^+^ TAMs in Colon Cancer

**DOI:** 10.1101/2025.09.02.673578

**Authors:** Zhihao Wei, Yan Dong, Ruiyang Zi, Taorui Liu, Yahan Fan, Rui Zhang, Yongning Shang, Haoran Jiang, Xiang Zhao, Jianjun Li

## Abstract

The role of non-monocyte-derived, in situ proliferating tumor associated macrophages (TAMs) in tumor progression has been underappreciated. Proliferative TAM subsets likely represent critical mediators in establishing the immunosuppressive microenvironment, given the susceptibility of TAMs to be hijacked by malignant cells for pro-tumorigenic functions. However, the regulatory networks governing their self-renewal circuitry and lineage bifurcation remained undefined. Combining single cell RNA sequencing of human and murine colon tumors, we identified CD168^+^ TAMs, a hyperproliferative subpopulation with FOXM1-dependent maintenance of replicative competence. Spatial multi-omics and functional validation revealed that this subset exhibited peritumoral localization and underwent immunosuppressive polarization through cholesterol-coordinated CX3CL1-CX3CR1 signaling. Colon cancer-specific lipid metabolic rewiring elevates glycerophospholipid and cholesterol flux, with PLA2G7-mediated glycerophospholipid catabolism promoting CX3CR1 expression in TAMs. Thus, the CX3CR1 signaling-driven immunosuppressive effect of CD168^+^ TAM is an integrated response to tumor-specific metabolic features. Targeting the CD168^+^ TAMs may enable therapeutic strategies for metabolic-immune reshaping in colon cancers.

## Introduction

Single-cell RNA sequencing (scRNA-seq) studies have identified MKI67^+^ tumor-associated macrophages (TAMs) within the tumor microenvironment (TME) across multiple cancer types^1,2^. This distinct subpopulation exhibits elevated expression of cell cycle-related genes, including *MKI67*, *TOP2A*, and *PCLAF*, suggesting inherent proliferative potential. While peripherally infiltrating monocytes are known to undergo TAM polarization under tumor microenvironmental cues, the paradoxical coexistence of diminished immune cell infiltration and persistent TAM expansion in advanced tumors implies that the role of self-renewal mechanisms in TAM population dynamics may have been underestimated^3^. The remarkable plastic adaptations of TAMs manifest as substantial phenotypic heterogeneity. Although canonical phenotypes such as SPP1^+^ and TREM2^+^ TAMs have been shown to contribute to establishing immunosuppressive niches, the absence of lineage-determining studies has left the gene regulatory networks (GRNs) governing their phenotypic evolution unresolved^4,5^. Therefore, deciphering the polarization dynamics of MKI67^+^ TAMs may offer novel insights into the mechanistic underpinnings of immunosuppressive TME formation.

Spatial organization and non-energy provisioning roles of lipid metabolism are critically implicated in TAM functionality. Spatial transcriptomic profiling of colorectal carcinoma (CRC) reveals tumor tissues compartmentalized into stromal-segregated microregions demarcated by cancer-associated fibroblasts (CAFs) and TAMs^6^. Conventional tumor microenvironment (TME) paradigms, long presumed to operate under homogeneous cellular distribution with unrestricted intercellular communication, are challenged by the discovery of macrophage-coated tumor clusters (MCTCs), underscoring the profound functional implications of TAM spatial positioning^7^. Lipid metabolic adaptations in TAMs exert functional impacts transcending mere energy provision. Dysregulated glycerophospholipid catabolism triggers endoplasmic reticulum (ER) stress that activates sterol regulatory cascades (e.g., SREBP signaling), establishing a pathological feedforward circuit driving aberrant cholesterol biosynthesis^8,9^. Cholesterol has been demonstrated to potentiate M2 polarization of TAMs within the CRC microenvironment^10^. Nevertheless, the cooperative mechanisms through which glycerophospholipid metabolism and cholesterol signaling orchestrate TAM lineage evolution under precise molecular subtyping conditions remain elusive.

This study interrogates the role of proliferative CD168^+^ TAMs in sculpting immunosuppressive colon cancer microenvironments through three interlocking dimensions: spatial compartmentalization, lineage trajectory, and metabolic adaptation. By integrating spatial transcriptomics with multiplexed immunofluorescence (mIF) mapping, we decode the topological organization and phenotypic features of CD168^+^ TAMs. scRNA-seq coupled with lineage tracing and functional perturbation models across *in vitro* and *in vivo* systems unravels their dynamic evolutionary vectors and molecular governance. Notably, we demonstrate that the immunosuppressive lineage polarization of CD168^+^ TAMs constitutes an adaptive response to the pathological lipid metabolic rewiring inherent to colon cancers.

### Transcriptional profiles and spatial organization of CD168^+^ TAMs

Integrated analysis of single-cell transcriptomes spanning 62 colorectal carcinomas (CRC) revealed a tumor-associated macrophage (TAM) subpopulation demonstrating proliferative competence through coordinated expression of mitotic regulators (including *MKI67*, *TOP2A*, *BIRC5*, and *PCLAF*; Figures 1A and 1B). This subset exhibited differential expression of the hyaluronan-sensing surface receptor *CD168* (*HMMR*) and transcription factor *FOXM1* (Figure 1B). Notably, pan-cancer conservation of this CD168^+^ TAM signature was validated across hepatocellular carcinomas (HCC) and pancreatic ductal adenocarcinomas (PDAC) (Figures S1A-D). Interrogation of The Cancer Genome Atlas (TCGA) transcriptomes demonstrated that elevated CD168^+^ TAM signature scores portended significantly reduced overall survival across colon adenocarcinoma (COAD), LIHC, and PAAD cohorts (Figures S1E-G).

**Figure 1.**
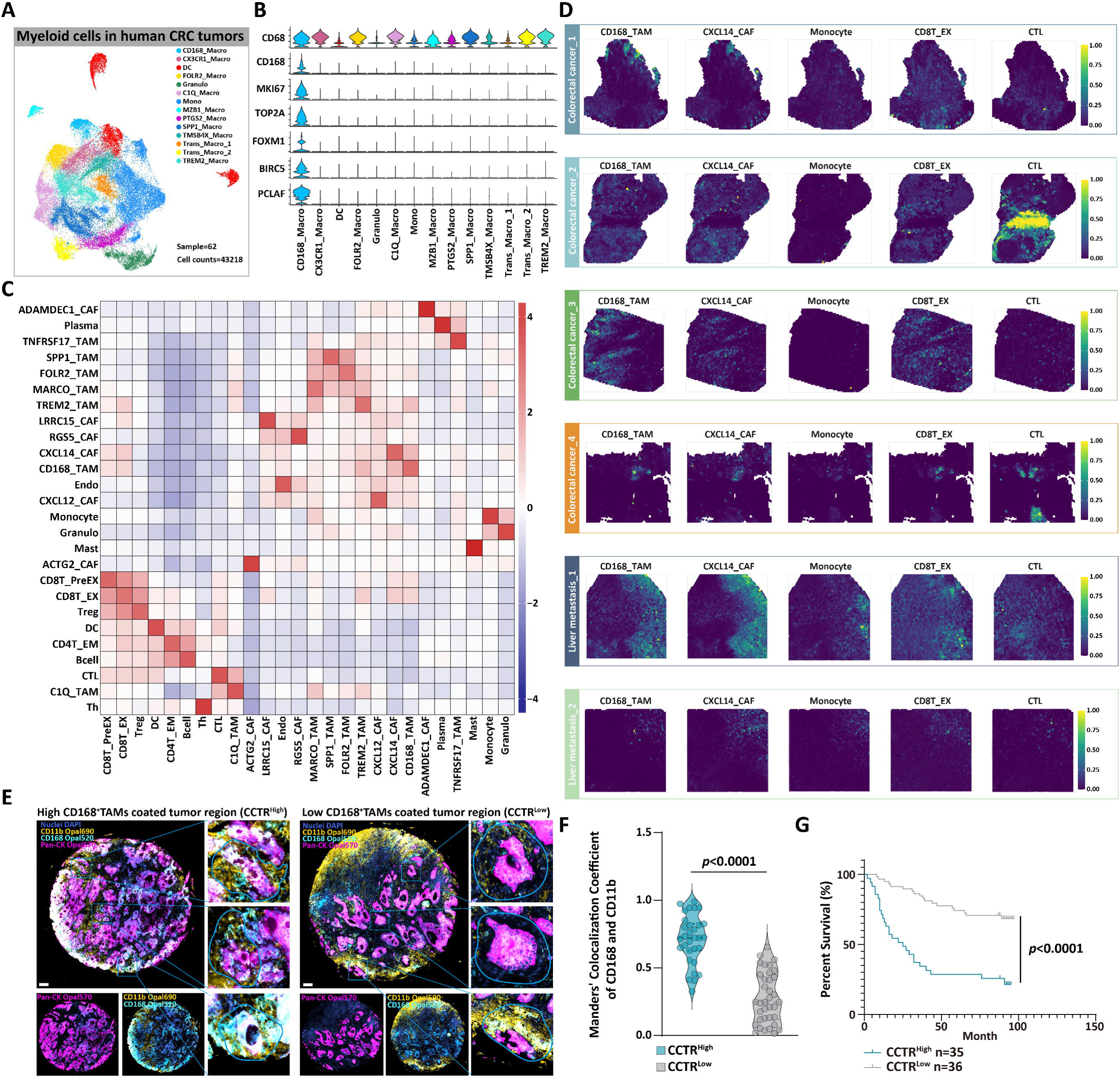
Transcriptional profiles and spatial organization of CD168^+^ TAMs. (A) Uniform Manifold Approximation and Projection (UMAP) of single-cell RNA sequencing (scRNA-seq) data (GEO: GSE178341) of myeloid cells (n=43218) isolated from CRC patients (n=62). (B) Single-cell transcriptomic profiling identified gene signatures of the CD168^+^ TAM subsets. (C) Heatmap showing the pairwise *Pearson* localization correlations between cell types in the spatial transcriptomic data (four primary colon cancer tissues and two hepatic metastases). (D) The spatial mapping of CARD-inferred illustrative cell types across samples. (E) Representative mIF images of pan-CK, CD11b, and CD168 in CRC patients. Three CCTR-defined regions of interest (ROIs) are highlighted with magnified views in each image. Scale bars = 200 μm. (F) CD168/CD11b co-localization analysis in CCTR^High^ versus CCTR^Low^ specimens reveals differential abundance of CD168^+^ TAMs, unpaired Student’s two-tailed t-test. (G) Kaplan-Meier overall survival curves of CCTR^High^ and CCTR^Low^ patients. *p* values were computed using Cox regression models.

Spatial compartmentalization fundamentally dictates cellular functionality and crosstalk within evolving tumor ecosystems. To decode the niche-specific behavior of CD168^+^ TAMs, we integrated spatial transcriptomics from four primary colon cancers and two hepatic metastases. Employing a conditional autoregressive deconvolution (CARD) framework informed by CRC scRNA-seq reference profiles, we resolved microenvironmental architecture at spot-level resolution^11^. Co-occurrence analyses revealed that TAMs and cancer-associated fibroblasts (CAFs), as well as T cells and DC cells, were cardinal colocalization modules (Figure 1C). CXCL14^+^ CAFs are pivotal in shaping the immunosuppressive CRC microenvironment^12^. Notably, among various TAM subtypes in primary and metastatic samples, CD168^+^ TAMs showed the strongest colocalization with CXCL14^+^ CAFs, coinciding with terminally exhausted CD8^+^ T cells (Figures 1C-D and S1h). Conversely, these macrophage subsets exhibited significant spatial exclusion from cytotoxic CD8^+^ T lymphocyte (CTL) infiltrates (Figures 1C and 1D). These findings suggest a potential immunosuppressive role for CD168^+^ TAMs.

Pan-cancer analyses reveal that macrophages are the predominant stromal cell populations that spatially organize with malignant cells to establish tumor microregions^6^. Employing multiplex immunofluorescence (mIF) on a 71-core colon cancer tissue microarray, we identified CD168^+^ TAMs forming distinct peritumoral architectures, which manifested as CCTRs (CD168^+^ TAM-coated tumor regions) (Figure 1E). Cases were stratified into CCTR^High^ and CCTR^Low^ cohorts using a median-driven quantitative threshold applied to CCTR areal coverage. Co-localization mapping demonstrated marked enrichment of CD168^+^ TAMs in CCTR^High^ cohorts (Figure 1F). Stratification-based survival analysis revealed that CCTR^High^ samples exhibited markedly inferior clinical outcomes compared to CCTR^Low^ counterparts (Figure 1G). These findings establish that CD168^+^ TAMs exhibiting proliferative competence drive the orchestration of immunosuppressive microregions to promote the progression of colon cancer.

### FOXM1 is essential for sustaining the CD168^+^ TAMs phenotype

We injected oncogene plasmids into the colonic region of mice to construct an orthotopic colon cancer model (Figure S3A and S3B). To investigate the functional dynamics of CD168^+^ TAMs, we harvested tumor tissues at three-time points (on the 14, 21, and 28 days) post-plasmid injection (Figure S3B). Flow cytometry was utilized to isolate CD11b cells for subsequent scRNA-seq analysis. Cross-species comparative analysis revealed conserved TAM subpopulations (Trem2, Folr2, Spp1, and Ptgs2 TAMs) in both human and murine colorectal carcinoma tissues (Figures 1A and S2A). Notably, integrated transcriptomic profiling identified FOXM1 as a transcription factor selectively enriched in CD168 TAM populations across species (Figures S2B and S2C). To investigate whether FOXM1 sustains the CD168 phenotype, macrophages were isolated and activated from healthy donor peripheral blood mononuclear cells (PBMCs). Functional validation through FOXM1 knockdown revealed a marked reduction in the CD168 population, as quantified by flow cytometry (Figures S2D and S2E).

The observed paucity of CD168^+^ TAM subsets within the tumor microenvironment may arise from FOXM1 downregulation during microenvironment-induced polarization, leading to diminished proliferative capacity. We found a time-dependent reduction in FOXM1 expression levels in CD168^+^ macrophages following co-culture with tumor cells (Figure S2F). CUT&Tag (Cleavage Under Targets & Tagmentation) profiling of FOXM1-binding sites in macrophages demonstrated selective enrichment at promoter regions of proliferation-associated genes (*MKI67*, *HMMR*, and *PCLAF*). Correspondingly, tumor cell co-culture induced marked attenuation of FOXM1 occupancy at these regulatory elements (Figures S2G–I). These findings establish FOXM1 as a key regulator of CD168^+^ phenotypic macrophages and reveal tumor cell-induced plasticity in this subset through microenvironmental reprogramming.

### The polarization dynamics of CD168^+^ TAMs

Our RNA velocity analysis of macrophages proximal to Cd168^+^ TAMs within the UMAP embedding revealed two divergent polarization trajectories for Cd168^+^ TAMs: Trem2 and Mmp9 lineage. And trajectory inference demonstrated that macrophages of alternative phenotypic states do not originate from Cd168^+^ TAM precursors (Figures 2A-B and S3C)^13^. During early tumorigenesis, Cd168^+^ TAMs predominantly committed to Trem2 lineage polarization, and temporal dynamics analysis revealed a progressive shift toward Mmp9 lineage with advancing tumor progression (Figure 2C). Strikingly, Mmp9^+^ TAMs still expressed Trem2, exhibiting a biphasic phenotype (Figure S3D). The risk of metastasis typically escalates with the tumor progression^14^. Comparative analysis of differentially expressed genes (DEGs) between the two lineages revealed that Trem2^+^ TAMs demonstrate enhanced lipid metabolic activity and marked immunosuppressive capacity. Conversely, Mmp9^+^ TAMs showed pronounced upregulation of pro-metastatic gene signatures and heightened microenvironmental adaptability (Figures 2D and S3E). These data suggest that Trem2^+^ TAMs promote an immunosuppressive microenvironment, while Mmp9^+^ TAMs facilitate tumor metastasis.

**Figure 2.**
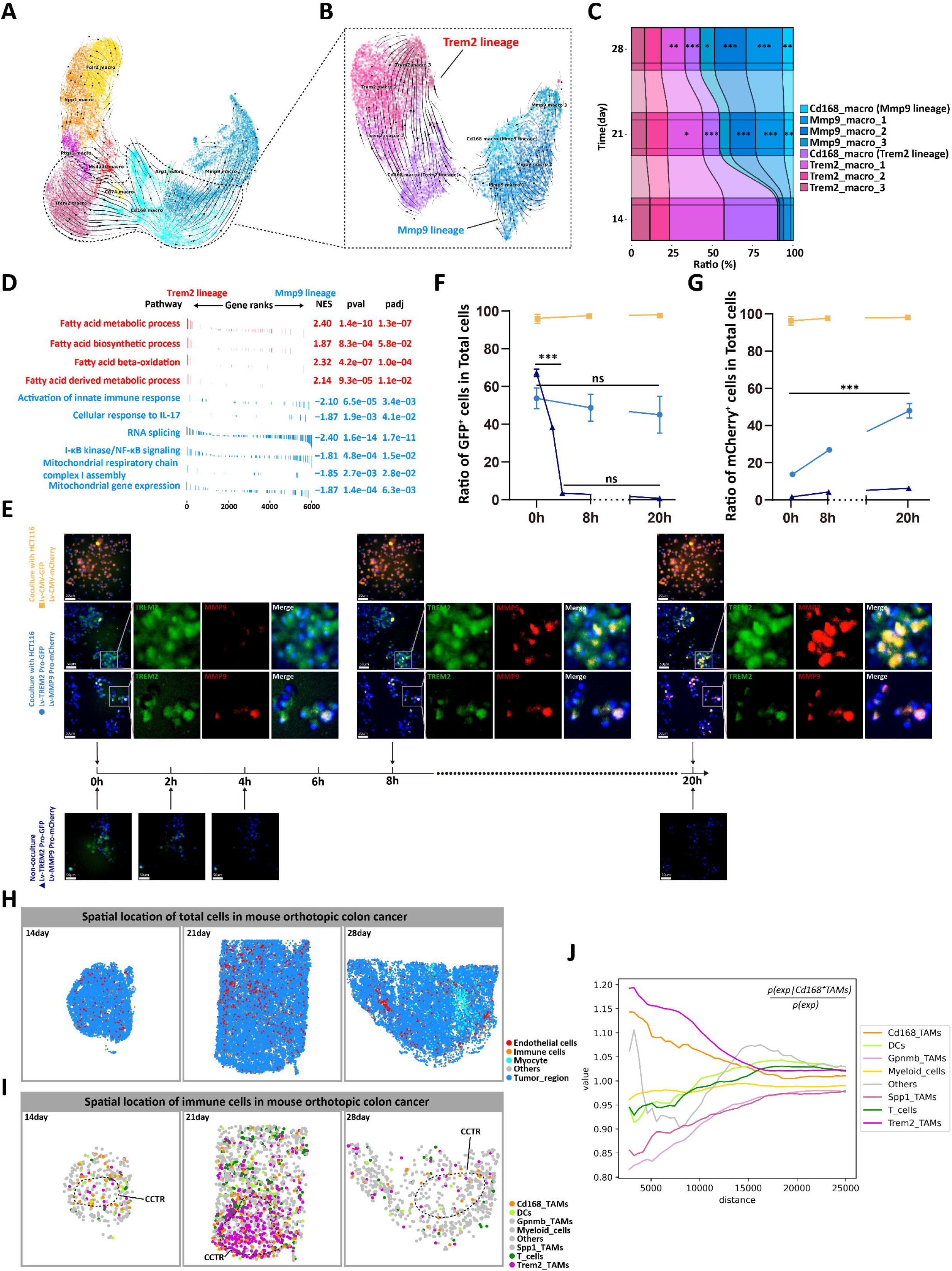
The polarization dynamics of CD168^+^ TAMs. (A) RNA velocity vectors of macrophages from mouse orthotopic colon cancer on UMAP embedding. (B) Polarization trajectories of Cd168^+^ TAMs along Trem2 and Mmp9 lineages. (C) Proportions from scRNA-seq data of Trem2 and Mmp9 lineage TAMs across the indicated time points. Pairwise comparison of subpopulation frequencies between late-stage (days 21/28) and early-stage (day 14) time points. Student’s two-tailed t-test, \**p*<0.05, \*\**p*<0.01, \*\*\**p*<0.001. (D) Gene set enrichment analysis (GSEA) on genes ranked by log2 fold change between Trem2^+^ TAMs and Mmp9^+^ TAMs. (E) Following isolation and activation of PBMC-derived macrophages, CD168^+^ cells were sorted and subjected to three experimental conditions: (i) Transduction with Lv-CMV-GFP and Lv-CMV-mCherry followed by HCT116 co-culture; (ii) Transduction with dual promoter-specific reporters (Lv-TREM2 promoter-GFP and Lv-MMP9 promoter-mCherry) with HCT116 co-culture; (iii) Dual promoter-specific reporter transduction without tumor cell co-culture. Live-cell imaging was conducted over 20 hours. The figures show representative images at the time points indicated by arrows. Scale bar = 50μm. (F) Frequencies of GFP^+^ cells among the total cells were statistically analyzed across different groups. One-way ANOVA, \*\*\**p*<0.001. (G) Frequencies of mCherry^+^ cells among the total cells were statistically analyzed across different groups. One-way ANOVA, \*\*\**p*<0.001. (H) Spatial mapping of total cells in the mouse orthotopic colon cancer. (I) Spatial mapping of immune cells in the mouse orthotopic colon cancer. (J) Proximity enrichment scores among cell subsets were calculated and visualized using the Squidpy method.

Subsequently, human PBMC-derived CD168^+^ macrophages were co-cultured with tumor cells in a Transwell system (Figure S3F). Real-time polarization dynamics of CD168^+^ macrophages were monitored via live-cell imaging. To functionally interrogate promoter activity, macrophages were transduced with dual lentiviral constructs harboring the TREM2 or MMP9 promoter regions, driving GFP or mCherry fluorescence reporters, respectively. Following co-culture with tumor cells at full confluency, macrophages underwent continuous live imaging over a 20-hour observation period to quantify promoter-driven fluorescent signals (Figure 2E). The results indicated that the transcription of TREM2 remained stable, while the transcription of MMP9 gradually intensified. The enhancement of MMP9 transcription occurred in both TREM2^+^ cells as well as in cells with initially weaker TREM2 transcription, which aligned with the biphasic nature of MMP9^+^ macrophages suggested by single-cell transcriptomics (Figure 2F-G). If not co-cultured with tumor cells, the transcription of TREM2 rapidly decreased within 4 hours, while the transcription of MMP9 remained consistently low. This *in vitro* experiment demonstrates that CD168^+^ macrophages initially undergo TREM2 lineage polarization in the tumor environment, with the MMP9 lineage polarization gradually increasing with the tumor advancement.

We employed spatial transcriptomics to investigate polarization dynamics of Cd168^+^ TAMs through spatially resolved assessment of murine colon cancer tissues harvested at sequential time points. Comparative transcriptomic analysis demonstrated that our *in situ* oncogene delivery model exhibits strong transcriptional concordance with Apc^KO^-induced colon tumors, characterized by elevated expression of *Lgr5*, *Axin2*, *Myc*, *Sox9*, *Prom1*, and *Lrig1*, alongside reduced levels of *Krt20*, *Muc2*, and *Cdx2* (Figures S3G and S3H)^15^. Spatial profiling of macrophage subpopulations defined by DEGs revealed that Trem2^+^ TAMs preferentially occupied hypovascular niches devoid of endothelial cells, whereas Spp1^+^ TAMs aggregated in vascular-rich territories (Figures 2H-I and S3J). This aligns with the results that Trem2^+^ phenotypes arise from proliferative Cd168^+^ TAMs polarizing under TME influences. CCTRs were identified within tumor tissues across three temporal phases (Figure 2I). Co-occurrence analysis showed Cd168^+^ TAMs had the strongest colocalization with Trem2^+^ TAMs, and they demonstrated reciprocal exclusion patterns with T cells and dendritic cells (DCs), suggesting immunosuppressive crosstalk within the TME (Figure 2J).

### CX3CR1 signaling drives the TREM2^+^ polarization in CD168^+^ TAMs

TREM2^+^ TAMs are critical in the establishment of immunosuppressive TME^4^. Application of RNA velocity analysis to scRNA-seq data derived from murine orthotopic colorectal cancer models revealed six genes—Cx3cr1, Il10ra, Apoe, Alox5, Fosb, and Cd36—exhibiting robust association with Trem2 lineage polarization of Cd168^+^ TAMs (Figure S4A). Elevated CX3CR1 expression, however, emerged as the sole prognostic biomarker significantly correlated with adverse clinical outcomes in colon cancer patients (Figure S4B)^16^. mIF analysis was performed on colon cancer tissue microarray to evaluate the spatial expression of CX3CR1 in CD168^+^ TAMs (Figure 3A). CX3CR1 expression exhibited a significant positive correlation with CD168 within both the tumor stromal region (Pan-CK^-^) and the macrophage population (CD11b^+^), demonstrating that CD168^+^ TAMs typically express CX3CR1 (Figures 3B and 3C). Advanced-stage tumor tissues exhibited a marked enrichment of CX3CR1^+^CD168^+^ TAMs compared to early-stage lesions (Figure 3D). Furthermore, Kaplan-Meier survival analysis revealed that elevated infiltration of CX3CR1^+^CD168^+^ TAMs within the TME was significantly associated with adverse clinical outcomes (Figure 3E).

**Figure 3.**
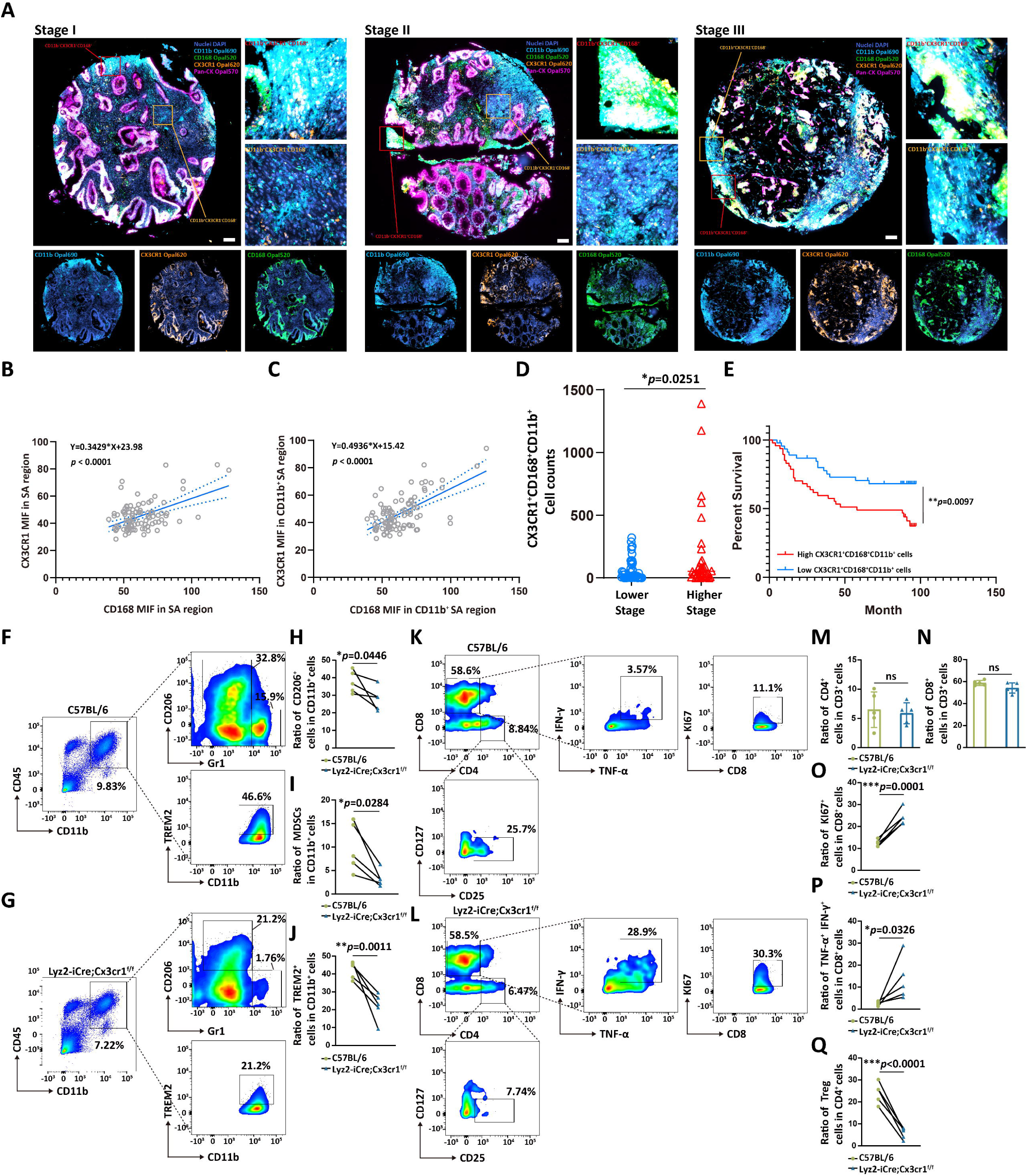
CX3CR1 activation promotes the generation of TREM2^+^ TAMs. (A) Representative mIF images of pan-CK, CD11b, CD168 and CX3CR1 in colon cancer patients. Magnified views highlight the CX3CR1^+^CD168^+^ macrophage populations (red-boxed regions) and CX3CR1^−^CD168^−^ macrophage subsets (yellow-boxed regions) within each experimental panel. Scale bar=200μm. (B) Correlation analysis of mean fluorescence intensity (MFI) between CD168 and CX3CR1 in tumor stromal regions across multiple specimens. (C) Correlation analysis of MFI between CD168 and CX3CR1 in macrophages across multiple specimens. (D) Samples were stratified into high-stage (beyond IIA) and low-stage (IIA or earlier) groups. The infiltration density of CX3CR1^+^CD168^+^ macrophages was compared between these cohorts, unpaired Student’s two-tailed t-test. (E) Samples were stratified into high-infiltration and low-infiltration groups based on the median level of CX3CR1^+^CD168^+^ macrophage, followed by Kaplan-Meier survival analysis between the two cohorts. *P-values* were computed using Cox regression models. (F-G) Contour plots of TAMs (CD45^+^CD11b^+^, left), M2-TAMs (CD11b^+^CD206^+^, upper right), MDSCs (CD11b^+^Gr1^+^, upper right) and TREM2^+^ TAMs (CD11b^+^TREM2^+^, lower right) from control (n=5) or Lyz2-iCre; Cx3cr1^f/f^ (n=5) mouse colon cancer tissues. (H-J) Frequencies of M2-TAMs, MDSCs, and Trem2^+^ TAMs in the tumor tissues of the two groups of mice, unpaired Student’s two-tailed t-test. (K-L) Contour plots of tumor reactive CD8^+^ T cells (CD8^+^INF-γ TNF-α Ki67, upper), Treg (CD4 CD25 CD127, lower) from control (n=5) or Lyz2-iCre; Cx3cr1^f/f^ (n=5) mouse colon cancer tissues. (M-N) Frequencies of CD4^+^ T and CD8^+^ T cells in the tumor tissues of the two groups of mice, unpaired Student’s two-tailed t-test. (O-Q) Frequencies of tumor reactive CD8^+^ T cells and Treg in the tumor tissues of the two groups of mice, unpaired Student’s two-tailed t-test.

Orthotopic colon cancers were established in myeloid-specific Cx3cr1 knockout mice (Lyz2-iCre; Cx3cr1^f/f^) to explore the role of CX3CR1 in inducing TREM2^+^ TAMs. Tumor tissues were harvested 28 days post-plasmid injection, and CD45^+^ cells were sorted for phenotypic analysis (Figure 3F-G and S4C-D). There was a marked decrease in Cx3cr1 expression within myeloid cells of Lyz2-iCre; Cx3cr1^f/f^ mice compared to the control C57BL/6 mice, while we observed no significant changes in the expression of CX3CR1 in T cells, B cells, and NK cells (Figure S4E). Conditional knockout of Cx3cr1 resulted in a marked reduction in infiltration levels of immunosuppressive cell populations, including M2 macrophages, myeloid-derived suppressor cells (MDSCs), and TREM2^+^ TAMs, within colon tumor tissues (Figures 3H-J and S4F). Concurrently, there was an enhanced infiltration of DC cells and T cells in the TME of Lyz2-iCre; Cx3cr1^f/f^ mice (Figure S4F). There was no difference in the proportion of CD4^+^ and CD8^+^T cells between the two groups (Figures 3M and 3N). However, in the Lyz2-iCre; Cx3cr1^f/f^ mice group, we observed higher expression of Ki67, TNFα, and IFNγ within CD8^+^ T cells, indicating a stronger tumor reactivity (Figures 3K, 3L, 3O, and 3P)^17^. Additionally, the infiltration of Tregs in the TME of the Cx3cr1 floxed group was reduced (Figure 3Q). In summary, the activation of CX3CR1 promotes the TREM2 lineage polarization of CD168^+^ TAMs, thereby exacerbating the immunosuppressive TME.

### MAF/MAFB orchestrate transcriptional programming of TREM2^+^ TAMs

To delineate the molecular underpinnings of TREM2 lineage polarization of CD168^+^ TAMs, we leveraged the CellOracle strategy to map transcriptional regulatory dynamics across discrete functional states of CD168^+^ TAMs^18^. CellOracle successfully identified and simulated the Trem2 and Mmp9 differentiation trajectories of Cd168^+^ TAMs using our scRNA-seq data of TAMs in mouse orthotopic colon cancer (Figures 4A and 4B). Computational screening identified Maf and Mafb as core transcription factors (TFs) within the Trem2 lineage. Subsequent in silico perturbation modeling of these regulators revealed state-transition suppression in Cd168^+^ TAM polarization trajectories, with Maf/Mafb knockout selectively inhibiting Trem2^+^ TAM specification while exhibiting limited perturbation on Mmp9^+^ TAM ontogeny (Figures 4C and 4D).

**Figure 4.**
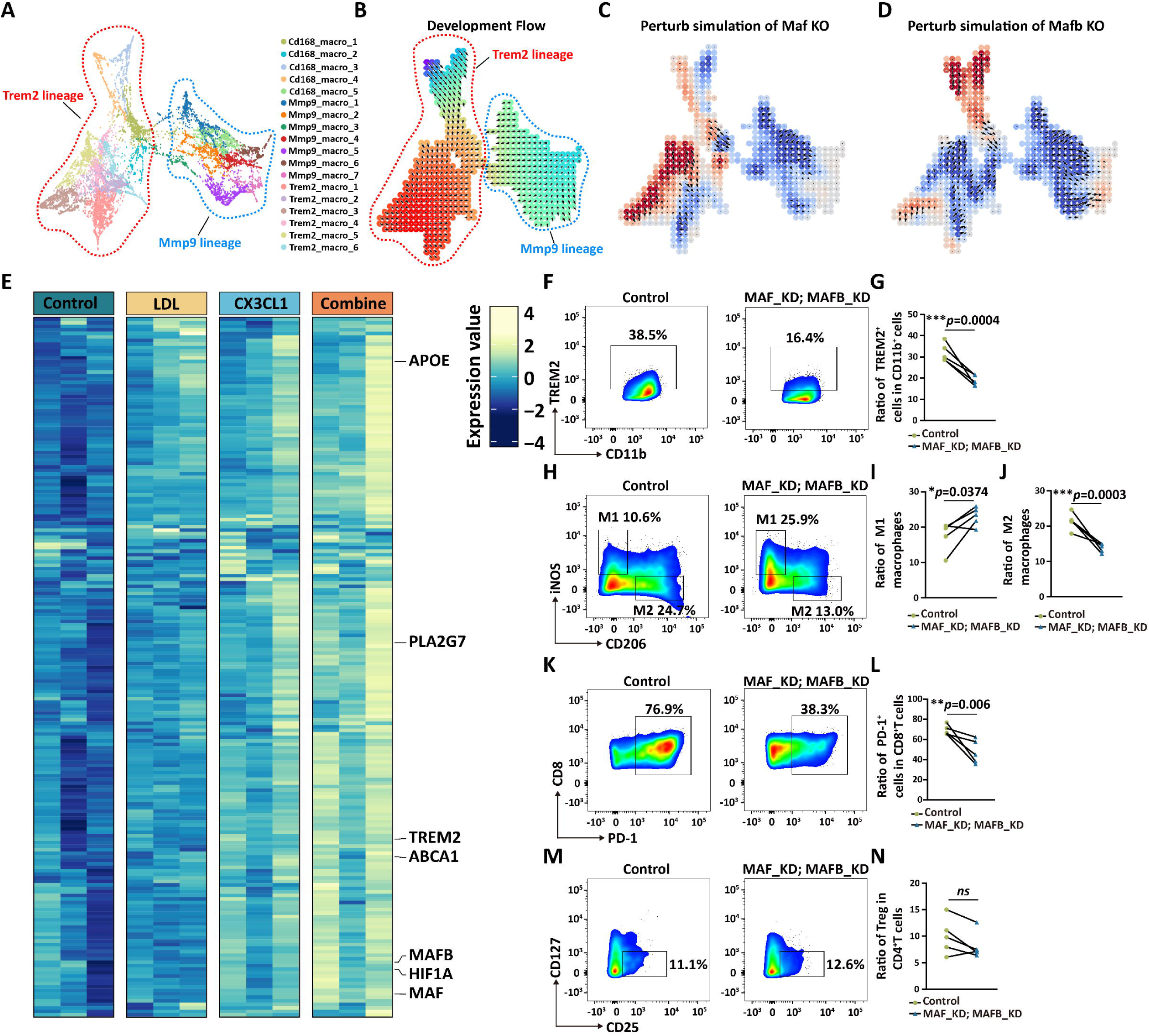
MAF/MAFB orchestrate transcriptional programming of TREM2^+^ TAMs. (A) Force-directed graph of Trem2 and Mmp9 lineage macrophages from scRNA-seq data of mouse orthotopic colon cancer. (B) CellOracle vector field graphic shows the developmental flow of Trem2 and Mmp9 lineage trajectory. Arrows start from Cd168^+^ TAMs and point towards the polarized subpopulation. (C-D) CellOracle KO simulation results for Maf (C) and Mafb (D). Perturbation scores were calculated to quantify the effect of TF knockout on the polarization of CD168^+^ TAMs, with negative values (blue) indicating suppression of normal polarization trajectories and positive values (red) reflecting enhancement of native polarization patterns. (E) Human PBMCs-derived CD168^+^ macrophages were co-cultured with HCT116 cells in a Transwell system for 48 hours following the illustrated treatment. Heatmap shows the relative expression of genes synergized by CX3CL1 plus cholesterol in CD168^+^ TAMs. (F-G) Contour plots and frequencies of TREM2^+^ TAMs (CD11b^+^TREM2^+^) in control versus MAF/MAFB-KD cohorts of humanized subcutaneous xenograft mouse models, unpaired Student’s two-tailed t-test. (H-J) Contour plots and frequencies of M1-type (CD11b^+^iNOS^+^) and M2-type (CD11b^+^CD206^+^) TAMs in control versus MAF/MAFB-KD cohorts of humanized subcutaneous xenograft mouse models, unpaired Student’s two-tailed t-test. (K-L) Contour plots and frequencies of PD-1^+^CD8^+^T cells in control versus MAF/MAFB-KD cohorts of humanized subcutaneous xenograft mouse models, unpaired Student’s two-tailed t-test. (M-N) Contour plots and frequencies of Tregs (CD4^+^CD25^+^CD127^-^) in control versus MAF/MAFB-KD cohorts of humanized subcutaneous xenograft mouse models, unpaired Student’s two-tailed t-test.

Cholesterol plays an essential role in modulating CX3CR1 functional activity^19^. Using a Transwell co-culture system, we demonstrated that cholesterol acts synergistically with CX3CL1 to activate CX3CR1 signaling. While cholesterol alone failed to induce CX3CR1 activation, co-stimulation with CX3CL1 significantly enhanced receptor activation, driving the TREM2^+^ phenotype polarization of CD168^+^ TAMs (Figures S5A-C). RNA-seq analyses identified some transcripts synergistically induced by CX3CL1 plus cholesterol in human CD168^+^ TAMs, which were representative among the driver genes of TREM2 lineage transition; these genes encoded for lipid metabolism (*APOE*, *PLA2G7*, and *ABCA1*) and TF (*MAF*, *MAFB*, and *HIF1A*) (Figure 4E).

CRISPR-based screening studies have identified DUSP18 as a key enzyme that promotes cholesterol synthesis in the TME of CRC and inhibits antitumor immunity by affecting CD8^+^ T cells^20^. scRNA-seq data from CRC tissues were stratified into DUSP18-high and DUSP18-low subsets using median expression thresholds, enabling comparative analysis of myeloid cell populations across both cohorts (Figure S5D). Tumor specimens with DUSP18^high^ expression displayed significant enrichment of TREM2^+^ TAMs, coupled with a concomitant reduction in MMP9^+^ TAM populations, indicative of a cholesterol-dependent myeloid polarization shift (Figures S5E and 5F). Comparative transcriptomic analysis of CD168^+^ TAMs between DUSP18^high^ and DUSP18^low^ cohorts revealed differential upregulation of MAF and MAFB in DUSP18^high^ specimens (Figure S5G). Integrated analysis of TCGA RNA-seq data from colon cancer patients indicated that MAF and MAFB expression positively correlated with the transcript levels of CX3CR1 and TREM2 (Figure S5H). These findings suggested that MAF and MAFB were potential downstream TFs of CX3CR1.

To verify the function of MAF and MAFB, we injected MAF and MAFB knockdown (KD) macrophages into humanized subcutaneous xenograft mice via the tail vein (Figure S5I). Tumor tissues were excised and subjected to flow cytometric analysis of immune cell populations. Macrophage-specific MAF/MAFB knockdown resulted in a marked reduction of TREM2^+^ TAMs compared to controls (Figures 4F and 4G). This was accompanied by a proportional decline in M2-polarized macrophages and a corresponding increase in M1-polarized subsets (Figures 4H-J). Within the T cell compartment, CD8^+^ T cells exhibited reduced PD-1 expression, a hallmark of T cell exhaustion, in knockdown cohorts (Figures 4K and 4L). In contrast, Treg infiltration remained statistically indistinguishable between groups (Figures 4M and 4N). In summary, our data demonstrate that the CX3CR1 signal promotes the TREM2 lineage polarization of CD168^+^ TAMs by activating MAF and MAFB.

### Glycerophospholipid metabolism promotes CX3CR1 upregulation in Tumor-associated macrophages

The role of lipid metabolic diversity in shaping TAM phenotypes remains to be elucidated^21^. To investigate lipid metabolic reprogramming within the TME, we divided the orthotopic colon cancer mouse model into two groups after injecting oncogenic plasmids: one maintained on standard chow and the other administered a high-fat diet regimen. Tumor specimens were processed for metabolomic profiling. Untargeted metabolomic analysis revealed a marked elevation of glycerophospholipid species within tumors from the high-fat diet cohort (Figure 5A). The CIBERSORT algorithm was employed to investigate potential associations between glycerophospholipid-metabolizing enzymes and immune cell populations in CRC specimens from the TCGA database, revealing a significant positive correlation between PLA2G7 expression levels and myeloid cell infiltration (Figure S6A)^22^. Integrated analysis of the TCGA and Genotype-Tissue Expression (GTEx) datasets revealed consistent upregulation of PLA2G7 in colorectal, pancreatic, and gastric carcinomas compared to adjacent normal tissues (Figure S6B). Analysis of scRNA-seq data from CRC specimens stratified by median PLA2G7 expression revealed enhanced myeloid infiltration and significantly elevated CX3CR1 expression levels in the high-expression cohort compared to low-expression counterparts (Figures 5B and S6C). Performing GO enrichment analysis on the DEGs of CX3CR1^+^ TAMs compared to other TAMs subpopulations, CX3CR1^+^ TAMs were enriched for pathways relating to the glycerophospholipid metabolic process, RNA splicing, and fibroblast proliferation (Figure 5C). While PLA2G7 levels showed no significant association with CD168^+^ TAMs, they demonstrated a positive correlation with TREM2^+^ TAMs and an inverse relationship with dendritic cell abundance (Figure S6D). These findings suggest PLA2G7-mediated glycerophospholipid metabolism may drive macrophage phenotypic polarization through CX3CR1-dependent mechanisms.

**Figure 5.**
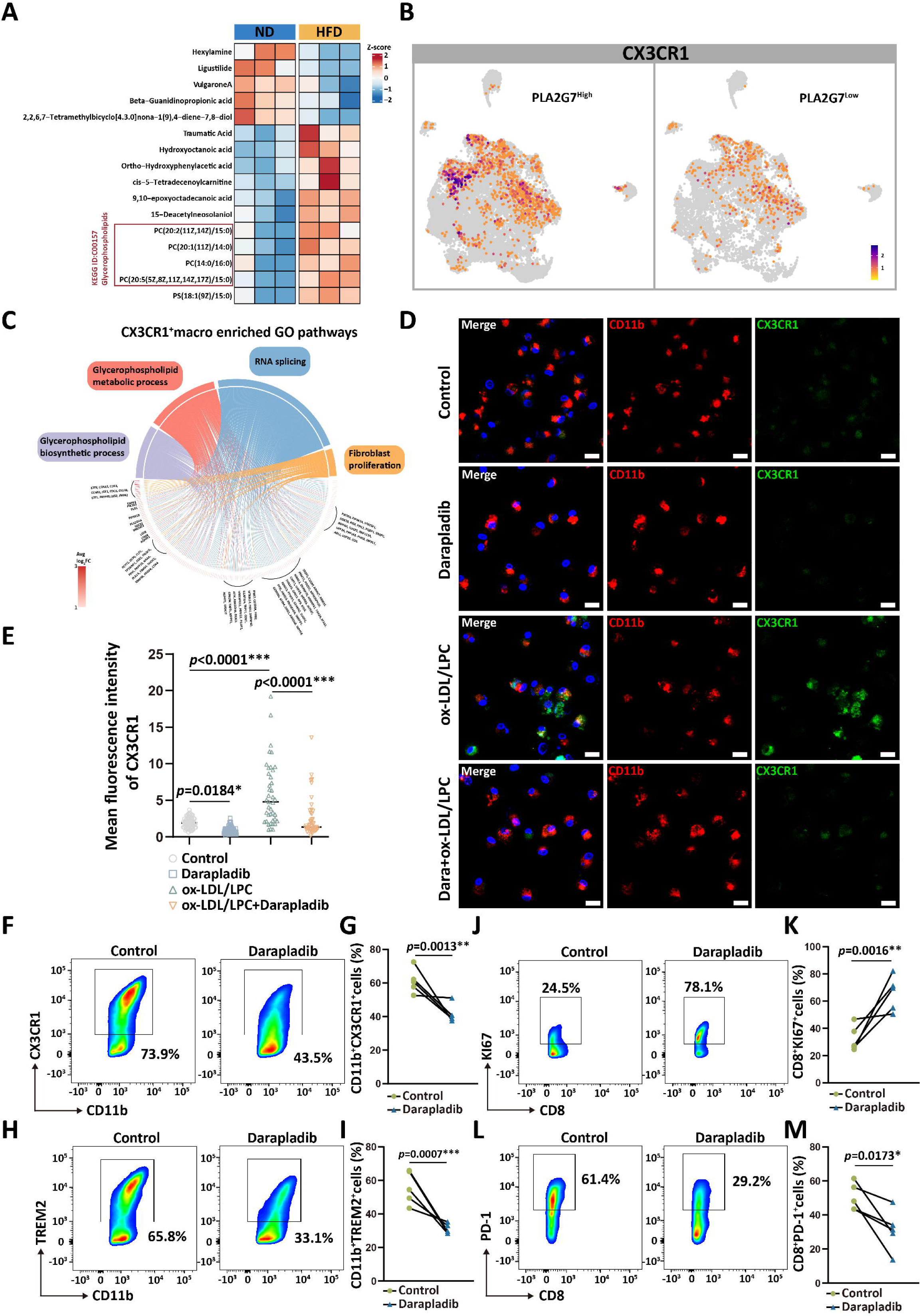
PLA2G7-mediated glycerophospholipid metabolism promotes CX3CR1 upregulation in TAMs. (A) Orthotopic colon tumors were harvested from mice fed a normal diet (ND) or a high-fat diet (HFD) at 21 days post-plasmid injection for untargeted metabolomic profiling. The heatmap depicts metabolites exhibiting significant differential abundance between the ND and HFD cohorts. (B) Comparative analysis of CX3CR1 expression in myeloid cells from PLA2G7^High^ versus PLA2G7^Low^ cohorts. (C) Chord diagram showing DEGs in CX3CR1^+^ TAMs enriched for GO pathways linked to glycerophospholipid metabolism, RNA splicing, and fibroblast proliferation. (D-E) PBMC-derived macrophages and HCT116 cells were co-cultured in a Transwell system for 48 hours under the illustrated conditions. CX3CR1 expression in macrophages was assessed by immunofluorescence staining. Scale bar=20μm. Quantitative analysis shows comparative mean fluorescence intensity of CX3CR1 on each cell across experimental groups, One-way ANOVA. (F-I) Mice received intraperitoneal administration of Darapladib (50 mg/kg) every other day, commencing 7 days post-intracolonic oncogene injection. Treatment continued until the experimental endpoint on day 21, when animals were euthanized for tumor tissue harvest. Orthotopic colon tumors of control (n=5) and Darapladib-treated (n=5) mice were collected and subjected to flow cytometry analysis. Contour plots and frequencies of CX3CR1^+^ (CD11b^+^CX3CR1^+^) and TREM2^+^ (CD11b^+^TREM2^+^) TAMs in control versus Darapladib-treated cohorts of mice with orthotopic colon cancers, unpaired Student’s two-tailed t-test. (J-M) Contour plots and frequencies of proliferative (CD8^+^Ki67^+^) and PD1^+^ (CD8^+^PD1^+^) CD8^+^ T cells in control versus Darapladib-treated cohorts of mice with orthotopic colon cancers, unpaired Student’s two-tailed t-test.

Lysophosphatidylcholine (LPC), a bioactive metabolite generated through phospholipase A2 group VII (PLA2G7)-mediated metabolism of oxidized low-density lipoprotein (ox-LDL), can be pharmacologically targeted by the specific PLA2G7 inhibitor Darapladib^23,24^. In a Transwell co-culture system modeling macrophage-tumor cell interactions, Darapladib treatment significantly attenuated CX3CR1 and TREM2 expression in macrophages. Conversely, exogenous supplementation with ox-LDL and LPC increased the prevalence of CX3CR1^+^ TAMs and TREM2^+^ TAMs compared to controls (Figures 5D-E and S7A-C). Mass cytometry (CyTOF) profiling of the tumor immune landscape in orthotopic colon carcinoma-bearing mice revealed diminished infiltration of immunosuppressive myeloid populations within the TME following Darapladib administration (Figures S7D-F). Corroborating the *in vitro* observations, phenotypic analysis of TAMs in mice orthotopic colon tumors demonstrated a marked reduction in CX3CR1^+^ and TREM2^+^ TAMs under Darapladib treatment compared to controls (Figures 5F-I). The remodeling of the immune microenvironment culminated in augmented tumor reactive capacity of CD8^+^ T lymphocytes coupled with concomitant decrease of terminally exhausted cells (Figures 5J-M and S7G-I). These findings establish that PLA2G7-driven glycerophospholipid metabolism within the TME governs CX3CR1 upregulation in TAMs.

### The therapeutic paradigm targeting CD168^+^ TAMs

Previous data have demonstrated that CD168^+^ TAMs contribute to the establishment of immunosuppressive TAM clusters through their proliferative capacity, peritumoral niche localization, and lineage evolution properties, with TREM2 lineage polarization emerging as a critical determinant of enhanced immunosuppressive functionality. Building on these findings, we conducted *in vivo* assays to evaluate the therapeutic potential of CX3CR1 signaling blockade in remodeling the immunosuppressive tumor microenvironment. The CX3CR1 blockade (JMS17-2, the specific inhibitor of CX3CR1) significantly attenuated orthotopic colon cancer progression and exhibited synergistic therapeutic efficacy when combined with PD-1 inhibition (Figure S8A). Spatial transcriptomic profiling of the tumor immune microenvironment revealed that JMS17-2 treatment reduced TREM2^+^ TAMs and significantly attenuated their co-localization with CD168^+^ TAMs, while showing no discernible impact on SPP1^+^ TAMs (Figures 6A-C and S8B-C). This effect was corroborated at the protein level through mIF analysis (Figures 6D-E). The observed effect concurrently elicited a marked elevation of CD8^+^ T cell populations in both neoplastic and paracancerous tissues (Figures 6D and S8D-E). These findings collectively demonstrated that CX3CR1 blockade substantially attenuated TREM2 lineage polarization of CD168^+^ TAMs.

**Figure 6.**
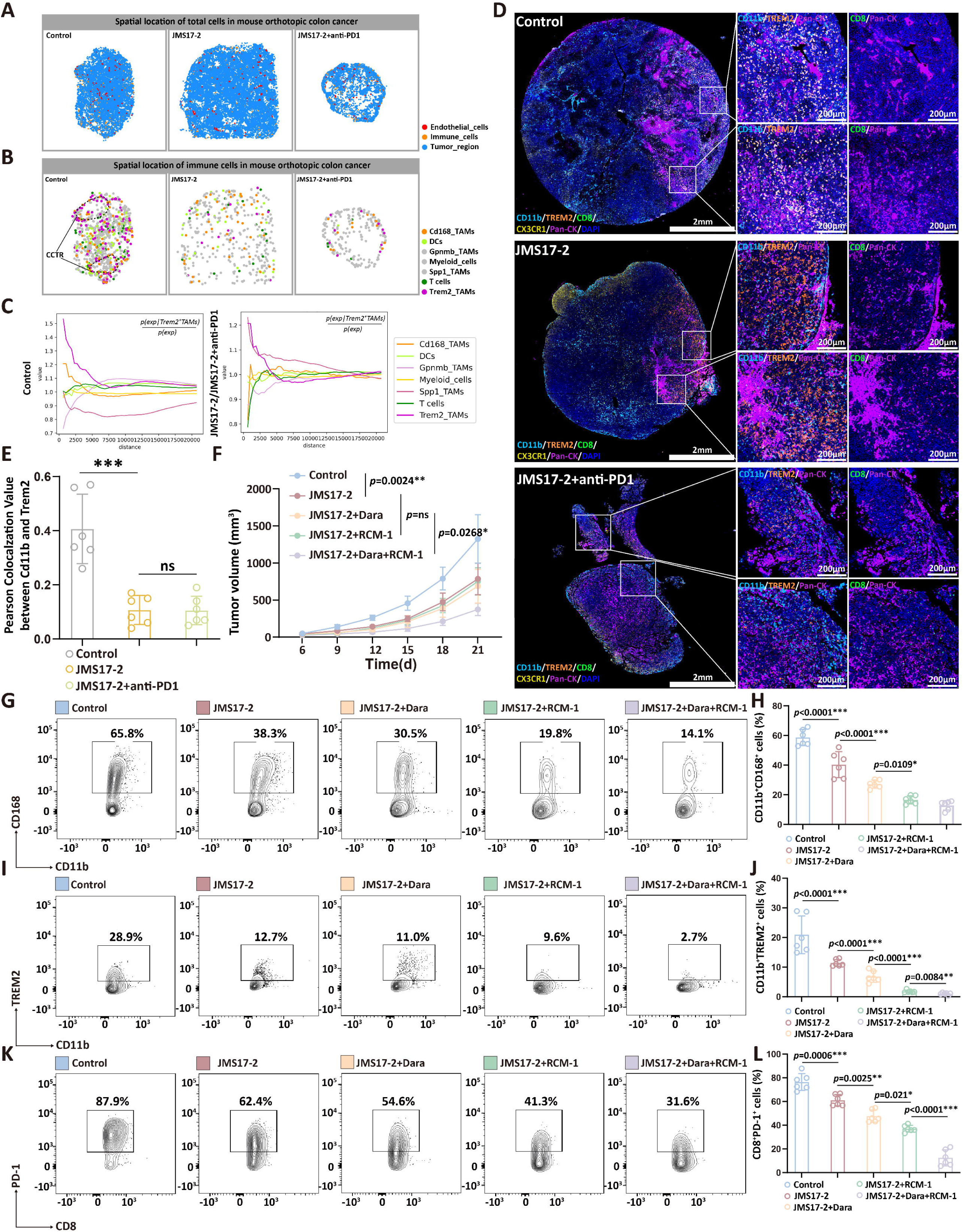
Validation of therapeutic strategies targeting CD168^+^ TAMs. (A) Spatial mapping of total cells in the mouse orthotopic colon cancer across different experimental groups. (B) Spatial mapping of immune cells in the mouse orthotopic colon cancer across different experimental groups. (C) Proximity enrichment scores between cell subpopulations were calculated and visualized using the Squidpy method. Left panel: Spatial enrichment patterns in control specimens. Right panel: Remodeled cellular interactions following JMS17-2 treatment (10mg/kg). (D) Representative mIF images of Pan-CK, CD11b, TREM2, CX3CR1, CD8 in mouse orthotopic colon cancer (Scale bar=2mm). The left magnified image indicates the abundance of TREM2^+^ TAMs in each experimental group by showing the colocalization of TREM2 and CD11b. The right magnified image shows the infiltration of CD8^+^ T cells in the corresponding region (Scale bar=200μm). (E) Each experimental group comprised three samples. Two representative fields of view from mIF images were analyzed for each sample. Pearson’s correlation coefficients for TREM2 and CD11b colocalization were quantified using the Coloc2 plugin in ImageJ. One-way ANOVA, \*\*\**p*<0.001. (F) Humanized mice bearing subcutaneous xenografts (n=6 per cohort) initiated treatment regimens as depicted 6 days post-engraftment. Tumor growth curves demonstrate longitudinal measurements (mm³) across experimental conditions. Data represent mean ± SEM, analyzed by two-way ANOVA with Tukey’s post-hoc test (\*\*\**P*<0.001, \*\**P*<0.01, \**P*<0.05). (G-H) The experimental endpoint was reached when any subcutaneous xenograft tumor exceeded 1,500 mm³. All mice were sacrificed, and the tumor tissues were collected and subjected to flow cytometry analysis. Contour plots and frequencies of CD168^+^ TAMs across experimental groups, one-way ANOVA. (I-J) Contour plots and frequencies of TREM2^+^ TAMs across experimental groups, one-way ANOVA. (K-L) Contour plots and frequencies of PD-1^+^CD8^+^ T cells across experimental groups, one-way ANOVA.

Next, we developed a dual-target therapeutic strategy against CD168^+^ TAMs, combining FOXM1 inhibition (RCM-1, a specific FOXM1 inhibitor) to suppress their proliferative capacity with concurrent blockade of the CX3CR1 signaling axis using JMS17-2 and Darapladib. This combinatorial approach demonstrated therapeutic efficacy in humanized subcutaneous xenograft mice (Figure S8F). Comprehensive immune profiling using flow cytometry revealed that RCM-1 treatment significantly reduced CD168^+^ TAMs (Figures 6G and 6H). Synergistic CX3CR1 blockade further diminished the TREM2^+^ TAM subpopulation, concomitant with a favorable remodeling of the tumor immune microenvironment (Figures 6I and 6J). This immunomodulatory shift was characterized by decreased expression of exhaustion markers (PD-1, TIM-3) on infiltrating CD8^+^ T lymphocytes (Figures 6K-L and S8G-H). Our findings demonstrate that targeting CD168^+^ TAMs represents a mechanistically grounded strategy to mitigate immunosuppressive circuitry in colon cancer.

## Discussion

Although TAMs can expand via local self-renewal in tumors, supported by scRNA-seq evidence of proliferative MKI67^+^ TAMs across malignancies, the functional identity and lineage trajectories of this subpopulation remained unresolved. Based on distinct membrane protein expression profiles, we defined this macrophage subpopulation as CD168^+^ TAMs. Spatial distribution analyses revealed that CD168^+^ TAMs exhibited a clustered organizational pattern in peritumoral regions, and FOXM1 emerged as the predominant transcriptional regulator governing their proliferative potential. Crucially, CD168^+^ TAMs bifurcated into immunosuppressive (TREM2 lineage) and pro-metastatic (MMP9 lineage) polarization within the TME. Mechanistically, cholesterol-coordinated CX3CL1-CX3CR1 signaling drove TREM2 lineage commitment through MAF/MAFB transcription factor activation. PLA2G7-mediated glycerophospholipid metabolism licensed CX3CR1 expression in TAMs, prompting our development of a dual therapeutic strategy combining PLA2G7 inhibition with CX3CR1 blockade. Preclinical models demonstrated that combinatorial targeting of FOXM1-dependent proliferation and CX3CR1 signaling disrupted CD168^+^ TAM expansion and decoupled TREM2 polarization, effectively reprogramming the immunosuppressive landscape in colon cancer.

The proliferative capacity of macrophages has been historically underappreciated in shaping pro-tumorigenic niches, yet emerging experimental evidence positions self-renewing macrophages as a pivotal source of TAMs^25,26^. FOXM1, a cell cycle-associated transcription factor orchestrating proliferation, metabolic reprogramming, and DNA damage repair^27^, has been predominantly studied in macrophages through the lens of polarization and migratory regulation^28–30^. Through integrated scRNA-seq of human and murine colon cancer tissues, we identified FOXM1 as a defining regulator in CD168^+^ TAMs, directly controlling the transcription of phenotype-specific genes (*MKI67*, *PCLAF*, and *HMMR*), thereby establishing its non-redundant role in sustaining TAM proliferative programs. While our current experiment data cannot fully resolve the spatial logic underlying CD168^+^ TAM encirclement of tumor clusters, their proliferative signature coupled with CX3CR1 expression suggests partial derivation from gut tissue-resident macrophages (TRMs)—a population known to undergo progressive peritumoral reorganization during tumorigenesis^31,32^. This developmental linkage may synergize with the TREM2 lineage polarization bias of CD168^+^ TAMs to mechanistically explain the predominance of TREM2^+^ TAMs in forming macrophage-coated tumor clusters (MCTCs)^7^.

While MAF and MAFB have been identified as master transcriptional regulators governing macrophage terminal differentiation^33^, with MAF specifically implicated in driving M2-like polarization^34^, the upstream mechanisms driving their activation remained elusive, hampering therapeutic targeting. Our work delineates that the cholesterol-coordinated CX3CL1-CX3CR1 signaling axis orchestrates TREM2 lineage commitment in CD168^+^ TAMs through MAF/MAFB activation. Although cholesterol is recognized as a cardinal metabolic determinant of immunosuppressive CRC microenvironments, its immunomodulatory mechanisms are mechanistically obscure^20^. CX3CR1 emerges as the sole chemokine receptor whose effector functions are directly modulated by cholesterol molecules^19^. Thus, our finding provides a definitive mechanism for cholesterol-mediated immunosuppression. Lipidomic profiling of clinical cohorts revealed elevated glycerophospholipid levels as a signature alteration in CRC^35^, a phenomenon validated in our high-fat diet-induced murine models through metabolomic analysis. Strikingly, PLA2G7-driven glycerophospholipid metabolism licenses CX3CR1 expression in macrophages, positioning CX3CR1 signaling as an integrative effector node responsive to CRC-specific metabolic rewiring. Collectively, our research highlights the role and mechanisms of proliferating CD168^+^ TAMs in orchestrating the immunosuppressive colon cancer microenvironment, offering novel therapeutic strategies for TAM-targeted treatment.

## Acknowledgements

This study was supported by grants from the National Natural Science Foundation of China (82573168, 82203669, 81672856, 82472382, 81803028), and The General Program of Chongqing Natural Science Foundation (CSTB2024NSCQ-MSX0524, CSTB2023NSCQ-MS X0596).

## Methods

Detailed methods are provided in the online version of this paper and include the following:

### KEY RESOURCES TABLE

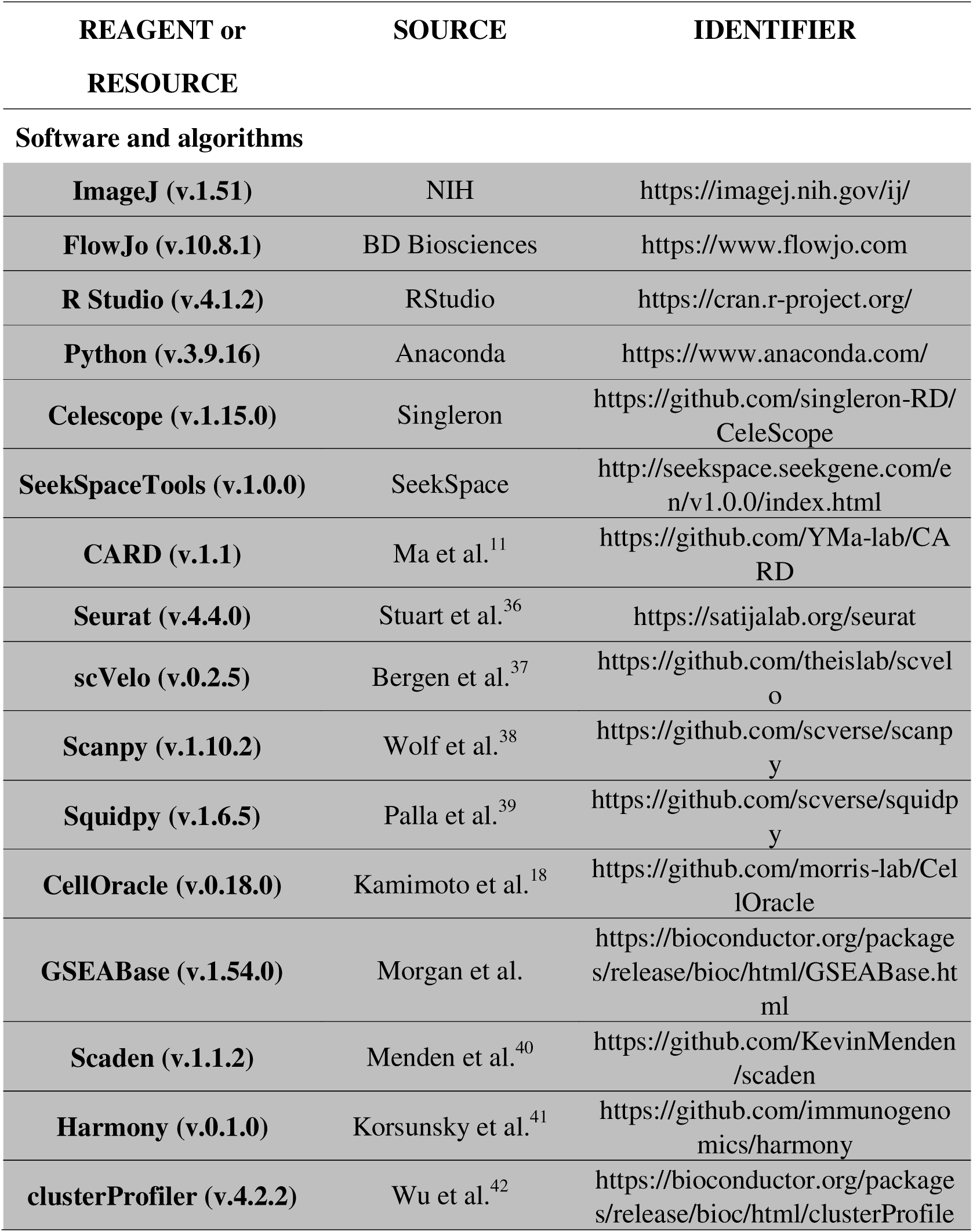
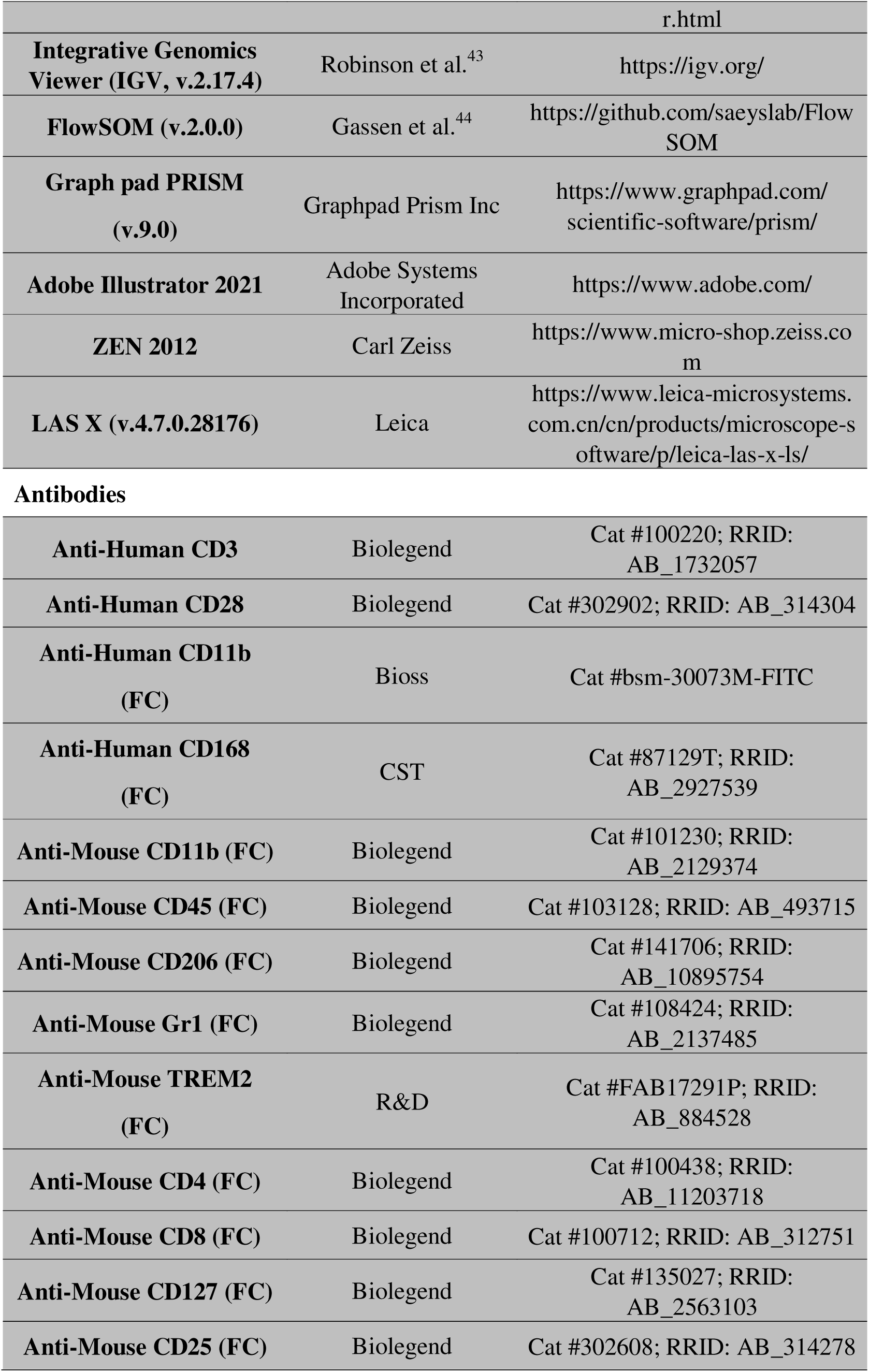
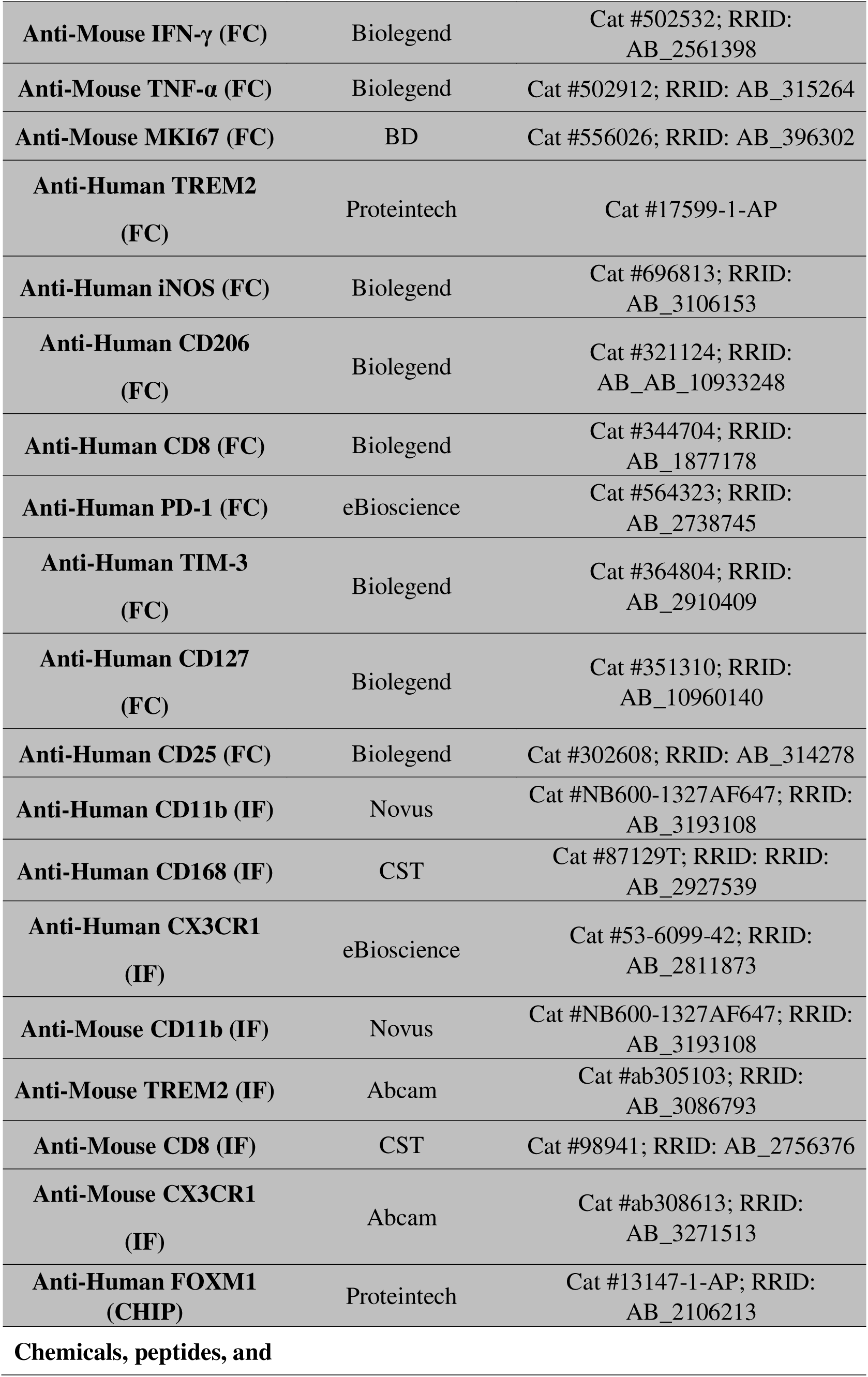
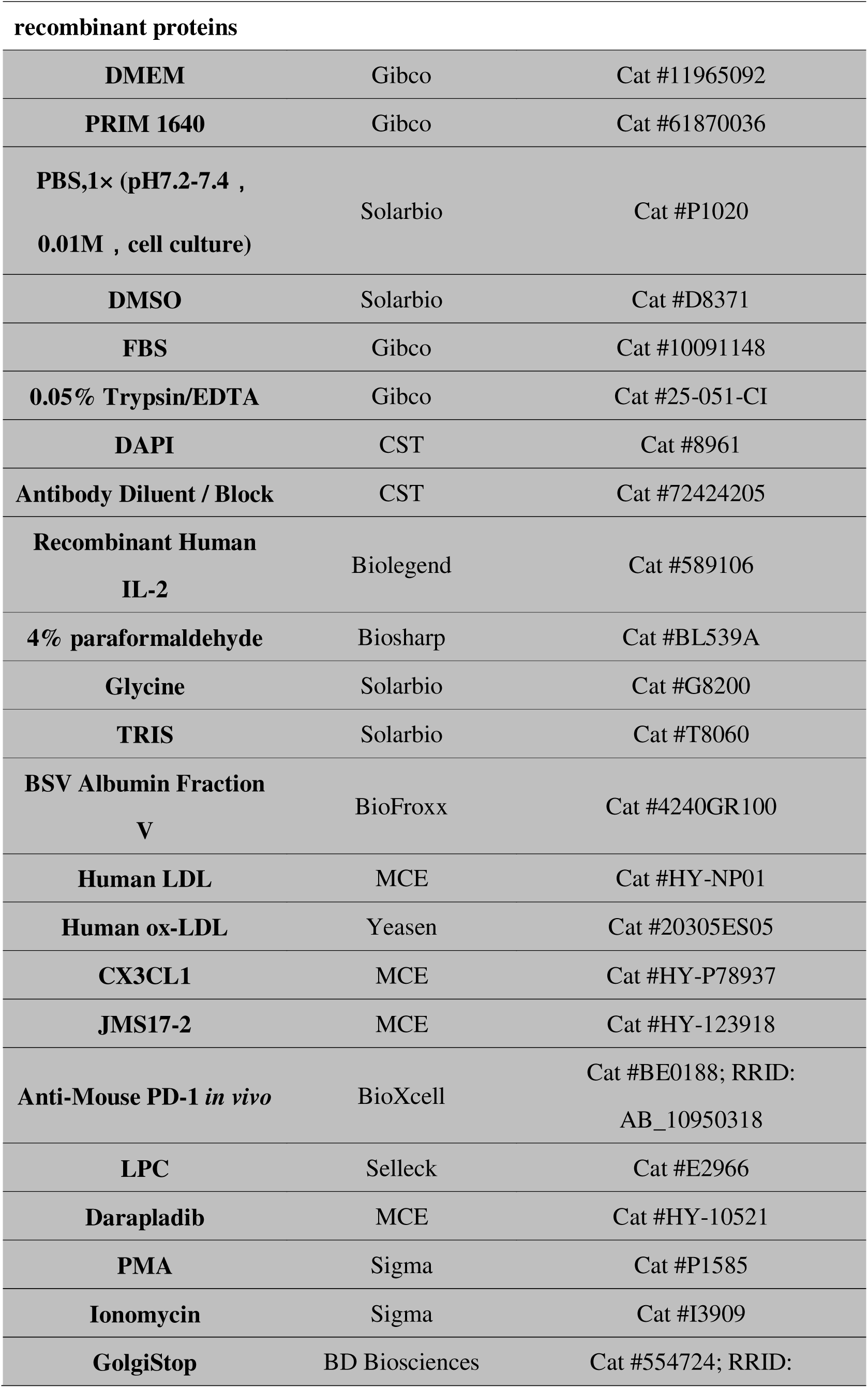
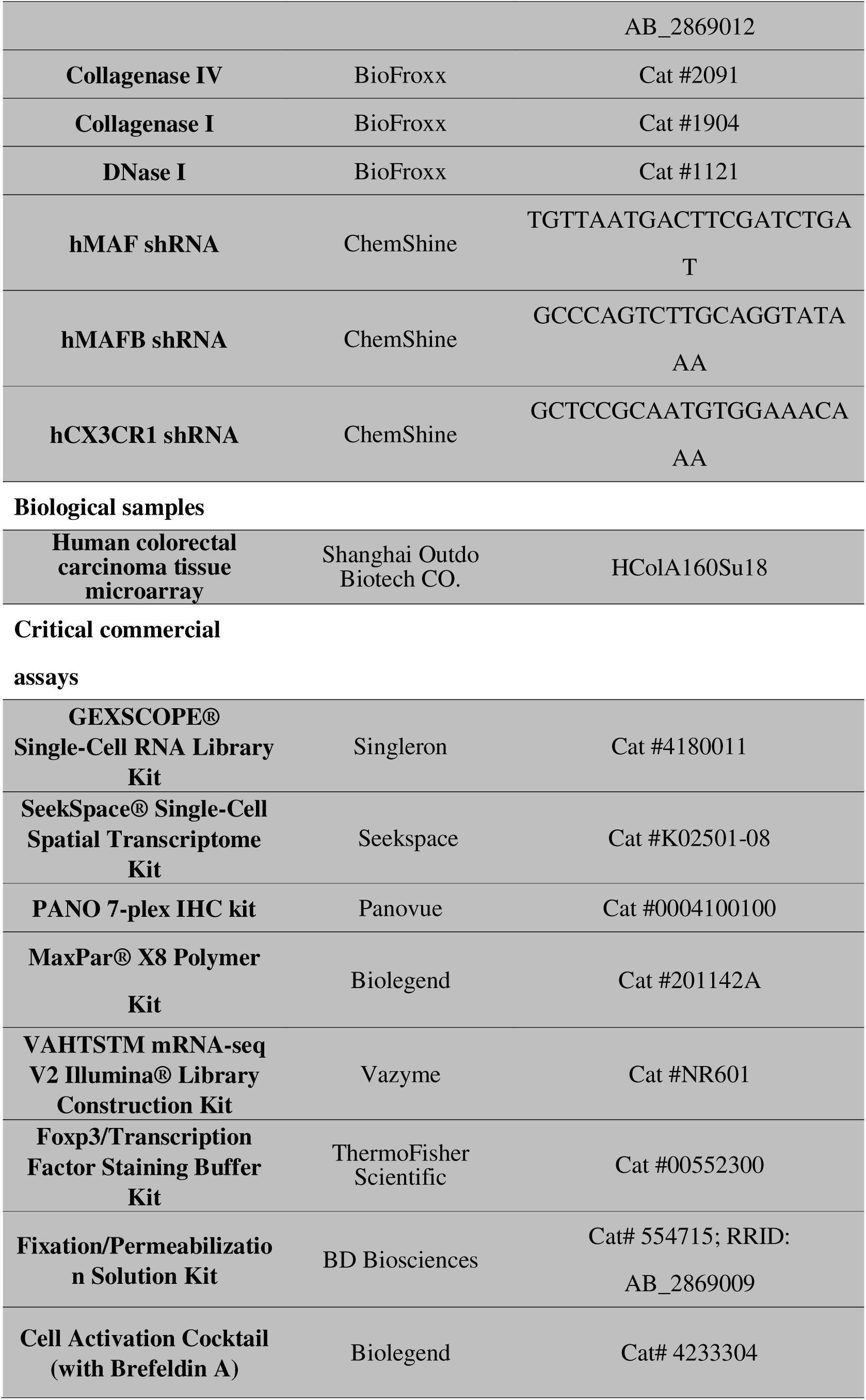
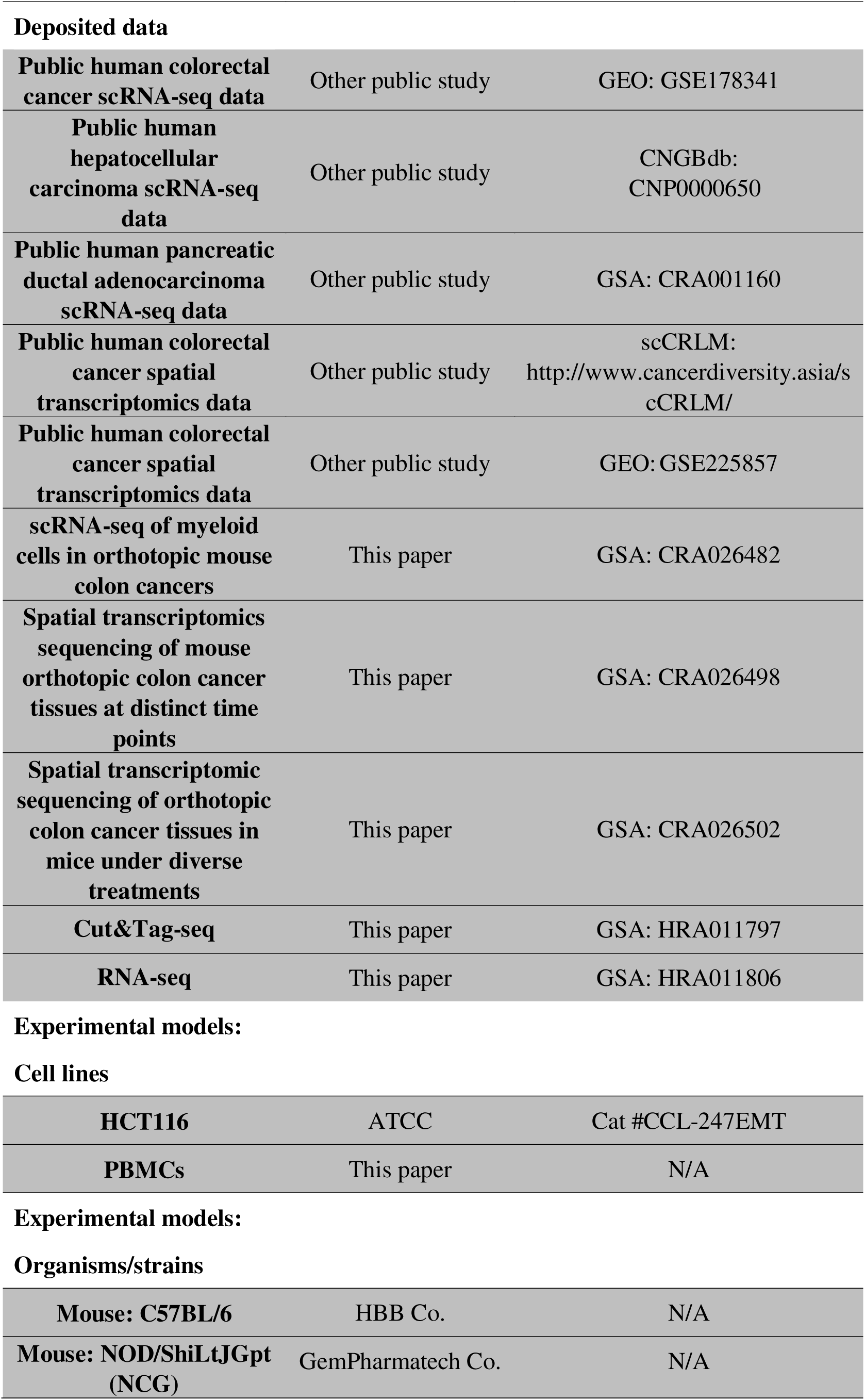

### RESOURCE AVAILABILITY

#### Lead contact

Further information and requests for resources and reagents should be directed to and will be fulfilled by the lead contact, Zhihao Wei.

#### Materials availability

This study did not generate new unique reagents.

#### Data and code availability

- The raw sequence data reported in this paper have been deposited in the Genome Sequence Archive (Genomics, Proteomics & Bioinformatics 2021) at the National Genomics Data Center (Nucleic Acids Res 2022), China National Center for Bioinformation / Beijing Institute of Genomics, Chinese Academy of Sciences. (GSA: CRA026482, CRA026498, and CRA026502; GSA-Human: HRA011806 and HRA011797) that are publicly accessible at https://ngdc.cncb.ac.cn/gsa-human. This paper has also analyzed existing, publicly available data. These accession numbers or relevant publications/database sources for the datasets are listed in the key resources table.
- No original code was generated for this study.

### EXPERIMENTAL MODEL AND SUBJECT DETAILS

#### Colon cancer tissue microarray

Colon cancer tissue microarray and patient clinical information (n = 93, Cat. No. HColA180Su18) were purchased from Shanghai Outdo Biotechnology Co. Ltd., Shanghai, China (https://www.superchip.com.cn).

#### Mouse orthotopic colon cancer model

Six-week-old male C57BL/6J mice were procured from VitalRiver Co., Ltd. Plasmids pT3-AKT (Cat #SZHY-0001), pT3-Myc (Cat #SZHY-0002), and pCMV-SB100 (Cat #SZHY-0003) were obtained from ShouzhengPharma Co., Ltd. A plasmid cocktail (AKT: Myc: SB100 = 1:1:0.2 μg ratio) was prepared in 5μL Ringer’s solution. Mice were anesthetized via intraperitoneal injection of tribromoethanol (250 mg/kg), positioned supine, and the abdominal fur disinfected with 70% ethanol. Following laparotomy, the colon was exteriorized using curved forceps with minimal traction to prevent tissue damage. The plasmid cocktail was injected into the serosal layer using an insulin syringe. Injection sites were compressed with sterile cotton swabs to confirm the absence of leakage. The colon was repositioned, and the abdominal wall closed via layered suturing. Mice were monitored daily for signs of distress (lethargy, abdominal distension), and moribund subjects were euthanized for necropsy to determine etiology.

#### Humanized xenograft mouse model

Humanized mice with an immunodeficient background were established via subcutaneous xenograft as previously described^45^. Male NOD/ShiLtJGptPrkdcem26Cd52Il2rgem26Cd22/Gpt (NCG) mice (age: 6 weeks) were obtained from GemPharmatech Co., Ltd. (Jiangsu, China) and acclimated to the facility for 1 week under specific pathogen-free conditions. Monocytes/macrophages isolated from PBMCs of healthy volunteers were administered via tail vein injection (dose: 1 × 10 cells per mouse). 1 week post-administration, 1 × 10 HCT116 cells were inoculated subcutaneously into the right inguinal region of each mouse. Commencing 1 week post-tumor inoculation, human T cells activated using CD3/CD28 antibodies and interleukin-2 (IL-2) were administered weekly via tail vein injection (dose: 1 × 10 cells per mouse) until the experimental endpoint.

#### Isolation and processing of PBMCs

PBMCs were isolated from Buffy coats obtained as byproducts of whole blood donations from healthy volunteers. The Buffy coat was diluted 1:1 with phosphate-buffered saline (PBS). The diluted blood was layered onto human peripheral blood lymphocyte separation medium (TBD, Tianjing, China) at a 1:2 ratio (diluted blood: separation medium). Centrifugation was performed at 700 × g for 15 minutes at room temperature (acceleration: 9; deceleration: 2). The mononuclear cell layer was collected, resuspended in PBS, and washed three times. Cells were subsequently resuspended in 5–10 mL of red blood cell (RBC) lysis buffer and agitated thoroughly for 5 minutes. Following centrifugation at 300 × g (room temperature), the RBC-depleted mononuclear cell layer was collected. Isolated PBMCs were resuspended in RPMI-1640 medium supplemented with 10% fetal bovine serum (FBS) and adjusted to a density of 1 × 10 cells/mL. Cells were plated and incubated overnight at 37°C under 5% CO . Adherent cells were stimulated with 20 ng/mL macrophage colony-stimulating factor (M-CSF) for macrophage differentiation and maintained for 7 days. Non-adherent cells were transferred to six-well plates pre-coated with 2 μg/mL anti-CD3 antibody. T cells were activated by supplementing the medium with 2.5 μg/mL anti-CD28 antibody and 1000 U/mL interleukin-2 (IL-2), followed by culture for 3 days. This study received approval from the Institutional Review Board of Southwest Hospital (#KY2023115).

#### Tumor cell lines

HCT116 cell line was purchased from ATCC, propagated, and passaged as adherent cell cultures. HCT116 cells were maintained in complete McCOY’s 5A (Gibco) medium containing 10% fetal bovine serum (FBS) and 100U/ml penicillin/streptomycin. HCT116 cells were maintained in adherent conditions at 37°C in a humidified atmosphere containing 5% CO_2_. The medium was replaced three times weekly, and the cells were passaged using 0.05% trypsin/EDTA (Corning) and preserved at early passages.

### METHOD DETAILS

#### Transcriptomic analysis with The Cancer Genome Atlas (TCGA) datasets

##### Data preparing

The transcriptomic and clinical data for colorectal cancer (COAD), liver hepatocellular carcinoma (LIHC), and pancreatic adenocarcinoma (PAAD) were retrieved using the GDC download Tool from the Cancer Genome Atlas (TCGA) database (https://portal.gdc.cancer.gov). All data were normalized by log2 transformation.

##### Survival analysis

Cellular abundance of CD168^+^ TAMs in TCGA cohorts was deconvolved using Scaden method trained on scRNA-seq data specific to each cancer type^40^. Patients were dichotomized into high- and low-CD168^+^ TAM subgroups via optimal abundance threshold determination using the *maxstat.test* function from the maxstat R package. Association between CD168^+^ TAM stratification and overall survival (OS) was assessed for each malignancy using Cox proportional-hazards regression models (survival R package), adjusted for clinical stage as a covariate. Kaplan-Meier survival curves were generated and plotted with the *survfit* function (survminer R package) to visualize survival disparities.

#### Multiplex Immunofluorescence Staining and Analysis

##### Tissue microarrays preparation

Resected murine tumor tissues were collected within 30 minutes of resection and fixed in 10% formalin for 48 hours. Specimens were subsequently dehydrated through a standard ethanol gradient series, cleared in xylene, and embedded in paraffin blocks. Sections (4 μm) were mounted on glass slides, heated at 70°C for 1 hour to enhance adhesion, deparaffinized in xylene, and rehydrated through a graded ethanol series.

##### Multiplex Immunofluorescence Staining

Tissue microarrays were stained using the PANO 7-plex IHC Kit (Cat# 0004100100, Panovue, Beijing, China) following standardized protocols. Sections were deparaffinized in xylene, rehydrated through a graded ethanol series, and rinsed in tap water before heat-induced antigen retrieval. Tissue regions were demarcated with a hydrophobic barrier pen, followed by protein blocking with antibody diluent/blocking buffer. All antigens were sequentially labeled via an iterative protocol comprising: primary antibody incubation (1 hour, room temperature), secondary antibody incubation, tyramide signal amplification (TSA), and microwave-assisted antigen retrieval in stripping buffer to remove antibody-TSA complexes before proceeding to the next marker. Subsequently, sections were incubated with Opal Polymer HRP (10 minutes, 37°C), developed with TSA reagents from the PANO kit, counterstained with DAPI (5 minutes), and mounted with anti-fade medium. Whole-slide imaging was performed using an Olympus VS200 MTL microscope (Olympus Deutschland GmbH).

##### Data analysis

Image analysis software Qupath was used to build the algorithm by checking, training, and confirming the steps. Single cells were identified and quantified algorithmically, with positive cellular/regional classification determined using objectively defined intensity threshold criteria. Macrophages were defined as CD11b^+^PanCK^-^ cells, with subtypes further classified based on specific marker expression profiles. Co-localization of markers within individual cells was quantified using Mander’s correlation coefficient computed from co-expression signal intensities.

#### Bulk RNA sequencing and processing

Total RNA was extracted from samples according to the instructions of the Sangon Biotech (China) Trizol Total RNA Extraction Kit (#B511311). The quality and concentration of RNA were determined using a Qubit® 2.0 Fluorometer (Invitrogen). One microgram of RNA from each sample was taken for library preparation. The VAHTSTM mRNA-seq V2 Illumina® Library Construction Kit was used to add index codes to mark the sample sequences. The AMPure XP system (Beckman Coulter) was used to purify the library fragments to select cDNA fragments with a length of 150–200 bp. Then, 3 μL of USER enzyme (NEB) was used to treat the samples at 37°C for 15 min and at 95°C for 5 min. PCR amplification was performed using Phusion High-Fidelity DNA Polymerase, universal PCR primers, and index (X) primers. Finally, the PCR products were purified using the AMPure XP system, and the library quality was assessed on the Agilent Bioanalyzer 2100 system. The library was also quantified and pooled. Paired-end sequencing was performed on the Illumina NovaSeq 6000. The quality of the sequencing data was evaluated using FastQC (v 0.11.2), and the original reads were filtered using Trimmomatic (v 0.36). The remaining clean data were used for subsequent analyses. Clean reads were aligned to the human reference genome (hg38) using HISAT2 (v 2.0) with default parameters. RSeQC (v 2.6.1) was used to calculate alignment statistics. Qualimap (v 2.2.1) was used to examine the consistency distribution and genomic structure. BEDTools (v 2.26.0) was used to analyze gene coverage ratios.

#### Generation and processing of scRNA-seq data

##### Data generation

An orthotopic colon cancer model was established in mice via intraluminal plasmid injection of oncogenic constructs as previously described. Tumor tissues were harvested at days 14, 21, and 28 post-injection (n=3 mice/timepoint). Single-cell suspensions were prepared by enzymatic dissociation, followed by flow cytometry cell sorting of Cd45^+^Cd11b^+^ leukocyte populations for scRNA-seq. Suspensions (2×10 cells/ml in HyClone PBS) were loaded onto microchips using the Singleron Matrix® Single-Cell Processing System. Barcoded magnetic beads were recovered, and captured mRNA was reverse-transcribed into cDNA, amplified via PCR, and fragmented for adapter ligation. scRNA-seq libraries were constructed strictly according to the GEXSCOPE® Single-Cell RNA Library Kit protocol (Singleron). Libraries were sequenced on an Illumina NovaSeq 6000 platform (150 bp paired-end reads).

##### Data processing

Raw FASTQ files were processed with Celescope (v1.15.0) using default parameters. Reads were aligned to the mouse mm10 reference genome. High-confidence reads containing valid barcodes and unique molecular identifiers (UMIs) were retained to construct a gene expression matrix quantifying UMI counts per cell and gene. The gene count matrix was imported into R and processed with Seurat (v4.2.0). During Seurat object creation, cells were filtered based on the following thresholds: >200 and <6,000 genes detected per cell, and ≤20% mitochondrial gene content. Gene expression data were normalized using *NormalizeData* and scaled via *ScaleData*. The top 2,000 highly variable genes were identified with *FindVariableFeatures* for principal component analysis (PCA). Batch effects across samples were mitigated using Harmony (v0.1.0) by integrating the first 30 PCA dimensions via *RunHarmony*. UMAP was subsequently applied for two-dimensional visualization of cellular clusters.

##### RNA velocity and single-cell trajectories

TAMs of murine colon cancer tissues across multiple timepoints were integrated for RNA velocity analysis^13^, with cross-sample batch effects corrected using Harmony (v0.1.0). Cell clusters were defined according to marker-based manual annotation. The Seurat object was converted to Scanpy-compatible format (v1.10.2) via SeuratDisk (v0.0.0.9019), enabling subsequent analyses in Python (v3.9.16). Spliced and unspliced transcripts were annotated by processing cell-barcode-sorted BAM files (Celescope output) through the Velocyto pipeline. RNA velocity vectors were computed using scVelo (v0.2.5) with dynamical modelling to infer splicing kinetics^37^. Highly variable genes were defined as differentially expressed genes within TAM subpopulations, further refined by filtering via *scvelo.pp.filter_genes* (min_shared_counts=20). RNA velocity estimation was performed using *scvelo.tl.velocity* under default parameters. Velocity streamplots were generated by projecting velocity values onto the UMAP embedding through *scvelo.tl.velocity_graph* and visualized via *scvelo.pl.velocity_embedding_stream*.

##### In silico perturbation analysis

Computational perturbation analysis was performed using CellOracle (v0.18.0) following the official tutorial (https://morris-lab.github.io/CellOracle.documentation/) to identify key transcription factors (TFs) driving TREM2 lineage polarization in CD168^+^ TAMs^18^. The base gene regulatory network (GRN) was constructed using curated TF-target pairs from murine macrophages in CistromeDB. scRNA-seq data of TAMs from murine colon cancer were downsampled to 3,000 cells and 3,000 highly variable genes for subsequent analysis. TAMs were re-normalized and re-clustered using Scanpy (v1.9.3), with cellular trajectories recalculated via the *pt.set_lineage* function. The key TFs of each group were identified by overlapping factors ranked highly across degree centrality, betweenness centrality, and eigenvector centrality. MAF and MAFB were specifically selected as TREM2 lineage drivers with minimal impact on MMP9 lineage polarization. *In silico* knockdown of MAF/MAFB was simulated by setting their expression levels to 0. CellOracle then predicted trajectory shifts in cellular states using the simulated gene expression alterations.

#### *In vitro* macrophage lineage differentiation profiling by reporter assay system

An inducible lentiviral reporter system was engineered to validate polarization trajectories of CD168^+^ macrophages. Lentiviral constructs were generated as follows: Lv-TREM2 Promoter-GFP and Lv-MMP9 Promoter-mCherry (Experimental reporters); Lv-CMV-GFP and Lv-CMV-mCherry (Control reporters).CD11b^+^CD168^+^ macrophages isolated from PBMC-derived monocytes (7-day M-CSF stimulation at 20 ng/mL) were sorted and assigned to three treatment groups: a) Co-infection with control reporters, followed by Transwell co-culture with HCT116 cells (100% confluency); b) Co-infection with experimental reporters, followed by Transwell co-culture with HCT116 cells (100% confluency); c) Experimental reporter co-infection with Transwell culture in the absence of tumor cells. Live-cell imaging was performed for 20 hours post-treatment in a controlled-environment chamber (Leica DMi8 microscope). Lineage commitment dynamics of CD168^+^ macrophages were quantified by fluorescence intensity analysis.

#### GSEA analysis

Gene Set Enrichment Analysis was executed via R-based clusterProfiler (v4.2.2) on normalized expression data. GSEA ranked genes by phenotype correlation, calculating Enrichment Scores (ES) and Normalized Enrichment Scores (NES) through 1,000 permutations. Significance was determined by a False Discovery Rate (FDR) < 0.25, with significant gene sets visualized using enrichplot.

#### *In vivo* experiments

All animal experiments were conducted as specified and approved by the Animal Ethics Committee of the Third Military Medical University. Drugs used *in vivo*, along with their doses and administration routes, are listed below: 1) Darapladib, intraperitoneal injection every other day at 50 mg/kg; 2) JMS17-2, intraperitoneal injection daily at 10 mg/kg; 3) anti-PD1, intraperitoneal injection every three days at 200 μg/mouse; 4) RCM-1, intraperitoneal injection every other day at 20 mg/kg. Control mice received intraperitoneal injections of the same volume of solvent. For mice with subcutaneous xenografts, tumor size was measured in two dimensions every three days using calipers. Tumor volume was calculated using the formula (L × W²)/2, where L represents length and W represents width. The experimental endpoint was reached when the *in vivo* measured tumor volume of any subject mouse exceeded 1500 mm^3^. Subsequently, all mice were humanely euthanized, and tumor samples were collected for subsequent analysis. Regarding colon orthotopic tumors, mice underwent abdominal ultrasound examination every other day starting from day 21 post-plasmid injection. Tumor volume was calculated using the formula (L × W²)/2. Mice were euthanized when tumor volume exceeded 800 mm^3^ or when they exhibited signs such as ascites, abdominal distension, reduced activity, and poor spirits. Tumor samples were then collected for further analysis.

#### Tumor tissue processing and single-cell suspension preparation

Single-cell suspensions were prepared from tumor tissues using the DSC-200 Dissociator (RWD Life Science) with species-specific enzymatic kits: the Mouse Tumor Dissociation Kit (for murine orthotopic colon cancers, # DHTE-5001) or Human Tumor Dissociation Kit (for humanized xenografts, DHTEH-2505), per manufacturer’s protocols. Enzymatic cocktails were reconstituted in 5 mL HBSS according to manufacturer-specified ratios. Minced tumor tissues were transferred into tissue processing tubes containing the enzymatic solution. Tubes were mounted on the Dissociator and processed using program *H_Tumor_Heater_1* (optimized for soft tumors). Cell suspensions were filtered through 70 μm strainers, collected in 50 mL conical tubes, and centrifuged at 500 × g for 5 min (supernatant discarded). Pellets were resuspended in 1 mL RBC Lysis Buffer, incubated on ice for 3 min, then quenched with 6 mL DMEM. Samples were re-centrifuged (500 × g, 5 min; supernatant discarded). Cells were resuspended in RPMI 1640, DMEM, or appropriate buffer to the desired concentrations for downstream applications.

#### Mass cytometry

##### CyTOF Sample Preparation

Pre-conjugated antibodies were sourced from Fluidigm, or purified antibodies were labeled in-house using the MaxPar® X8 Polymer Kit (Fluidigm) per the manufacturer’s protocols. Metal-tagged antibodies were standardized to 200 μg/ml in Candor Antibody Stabilizer (Sigma-Aldrich) and titrated for optimal concentrations. Cells were washed in PBS, stained with 1 μM cisplatin (5 min, RT) for viability discrimination, then incubated with surface marker antibodies in staining buffer (PBS+0.5% BSA+0.02% NaN3) for 30 min at 4°C. After fixation, samples were incubated overnight with Cell-ID™ Intercalator-Ir (Fluidigm). Intracellular staining were performed using the Foxp3/Transcription Factor Staining Buffer Kit (eBioscience™) (30 min, 4°C), followed by washing and storage at 4°C until acquisition.

##### CyTOF Data Acquisition

Cells were washed twice in deionized water and spiked with EQ™ Four Element Calibration Beads (Ce140/Eu151/Eu153/Ho165/Lu175, Fluidigm). Samples were acquired on a Helios™ CyTOF instrument at event rates <300 events/s. Normalization and randomization of near-zero signals were performed using Helios™ software.

##### CyTOF Data Analysis

Viable singlets were gated manually. Batch effects were corrected via bead-based normalization. Immune cell subpopulations were resolved using FlowSOM clustering (R package *FlowSOM*, v2.0.0) applied to concatenated events from all samples.

#### Cut&Tag sequencing and data analysis

##### Data generation

Following cell viability assessment and counting via LUNA-FL, cells were conjugated to Concanavalin A-coated magnetic beads. Cell membranes were permeabilized using Digitonin. Under the guidance of a FOXM1-specific antibody, enzyme pA-Tn5 Transposase was directed to precisely bind and cleave DNA sequences proximal to the target protein, enabling factor-targeted tagmentation. During cleavage, adapter sequences were simultaneously ligated to the DNA ends. Adapter-ligated DNA fragments were amplified by PCR to generate the sequencing library. PCR products were purified using AMPure beads, and library quality was assessed on an Agilent Bioanalyzer 2100 system. Indexed samples were clustered on a cBot Cluster Generation System using the TruSeq PE Cluster Kit v3-cBot-HS (Illumina) according to the manufacturer’s protocol. Paired-end sequencing (150 bp) was performed on an Illumina NovaSeq 6000 platform by Sangon Biotech (Shanghai) Co., Ltd.

##### Data analysis

Raw paired-end reads in fastq format were preprocessed using fastp (v0.20.0)^46^ to remove adapter-containing reads, poly-N reads, and low-quality reads, yielding clean data. Q20, Q30 scores, and GC content were calculated from clean reads. All downstream analyses utilized these high-quality reads. Clean reads were aligned to the human reference genome (hg38) using BWA mem -k 32 -T 30 -t 4 -M (v0.7.12). Alignments were filtered to retain only high-quality (MAPQ ≥ 13), properly paired, non-PCR duplicate reads. Unique alignments were used for subsequent analysis. Genomic peak profiles were visualized using the Integrative Genomics Viewer (IGV, v2.17.4).

#### Untargeted Metabolomics Analysis

Resected mouse orthotopic colon tumor tissues were collected. For each sample, 25 mg of tissue was homogenized in 500μL extraction solvent (methanol/acetonitrile/water, 2:2:1 (v/v), containing an isotope-labeled internal standard mixture) using a tissue grinder at 35 Hz for 4 min. Samples were subsequently sonicated for 5 min in an ice-water bath. This homogenization-sonication cycle was repeated twice. Following extraction, samples were incubated at -40°C for 1 h and centrifuged at 12,000 rpm for 15 min at 4°C. The supernatant was transferred to injection vials for LC-MS analysis. Chromatographic separation was performed using a Vanquish UHPLC system (Thermo Fisher Scientific), coupled to an Orbitrap Exploris 120 mass spectrometer (Thermo Fisher Scientific). Raw data files were converted to mzXML format using ProteoWizard, followed by peak detection, extraction, alignment, and integration. Metabolite annotation was achieved by matching MS/MS spectra against the BiotreeDB (V2.1) library with an algorithm score cutoff set at 0.3.

#### Flow cytometry

Viability staining was performed using the Zombie NIR™ Fixable Viability Kit (BioLegend) per the manufacturer’s instructions to discriminate live/dead cells. Subsequent surface marker staining was conducted in FACS buffer at 4°C. Intracellular staining for nuclear proteins employed the eBioscience™ FoxP3/Transcription Factor Staining Buffer Set (Thermo Fisher Scientific), while cytoplasmic proteins (e.g., cytokines) were stained using the BD Cytofix/Cytoperm™ Fixation/Permeabilization Solution Kit (BD Biosciences). For intracellular cytokine detection, cells were stimulated with Cell Activation Cocktail (containing brefeldin A, BioLegend) for 4 hours at 37°C prior to staining. All samples were acquired on a BD FACS Aria™ III flow cytometer and analyzed with FlowJo software (v10.8.1).

#### Western blotting

Cells were lysed with an ice-cold protein lysis solution containing protease inhibitors (PMSF). Total protein concentration was measured by the BCA Protein Assay Kit according to the manufacturer’s instructions. Protein was loaded 10-30ug equally for separating by SDS-PAGE and transferred to nitrocellulose membranes (0.45μm). After blocking with TBST containing 5% BSA, membranes were incubated with primary antibodies and secondary antibodies. Blots were visualized with Affinity®ECL Reagent (femtogram).

#### Confocal microscopy

Cells were seeded on the confocal dish. Cells were fixed using 4% PFA and permeabilized using 3% Triton. After permeabilization, cells were blocked with 1% BSA in PBS for 30 min and incubated with primary antibody overnight at 4°C. Then, probed with Alexa 488-conjugated rabbit anti-mouse or Alexa 555-conjugated mouse anti-rabbit secondary. Finally, staining nuclei using DAPI. An LSM900 fluorescence confocal microscope (Zeiss) was used to obtain confocal fluorescence images.

#### Spatial transcriptome data generation and processing

Mouse colon cancer tissue samples were embedded in OCT compound (SAKURA, 4583). Sections (10 μm thickness) were prepared and transferred onto SeekSpace® chips, ensuring flat positioning without folds. Chips were mounted in SeekSpace® sc-Spatial Chip Holders, followed by assembly of the Space Chamber. Spatial barcoding was performed by adding 150 μl labeling reagent to the chamber, enabling transcript capture with location-specific barcodes. Sections were fixed and imaged by fluorescence microscopy. Tissue homogenization was conducted in pre-chilled lysis buffer using a Dounce homogenizer (KIMBLE, #D8938). Single-cell RNA sequencing (scRNA-seq) and spatial barcode libraries were prepared using the SeekSpace® Single-Cell Spatial Transcriptome Kit (K02501-08). Reverse transcription was first performed on the cells. An appropriate number of cells were then mixed with the ligation reagent and loaded into the sample well of the SeekOne® DD Chip S3 (Chip S3). Barcoded hydrogel beads (BHBs) and partitioning oil were added to their respective wells in Chip S3. Using the SeekOne® digital droplet system, the cell-laden ligation reagent and BHBs were co-encapsulated into water-in-oil emulsion droplets.

Emulsions were transferred to PCR tubes and incubated at 20°C for 60 min, followed by heat inactivation at 65°C for 10 min to generate barcoded cDNA and spatial barcodes. Decrosslinking was performed within droplets to recover barcoded cDNA and spatial barcodes. Sample indices were incorporated into pre-amplified spatial barcode products via PCR. Purified cDNA (20 ng) was used for index PCR amplification. Indexed libraries were purified with VAHTS DNA Clean Beads (Vazyme, N411), quantified using Qubit (Thermo Fisher Scientific, Q33226), and fragment size distribution assessed via Bio-Fragment Analyzer (Bioptic, Qsep400). Both scRNA-seq and spatial barcode libraries were sequenced on the Illumina NovaSeq 6000 platform using 150 bp paired-end (PE150) reads. FASTQ-formatted sequencing reads were aligned to the mouse reference genome assembly mm10. Gene expression matrices were generated using SeekSpace®Tools (v1.0.0). Spatial transcriptomics data, comprising the expression matrix, sample information, and spatial coordinates, were imported into the R environment using the CreateSeuratObject function to construct a Seurat object. Quality control (QC) criteria identical to those applied for scRNA-seq data were employed.

#### Analysis of spatial transcriptomics data

##### Cell type deconvolution and spatial co-localization analysis of public spatial transcriptomic data

To investigate the spatial localization of heterogeneous cellular populations in colorectal cancer tissues, we integrated 10x Visium data with scRNA-seq data using the CARD software package^11^. Briefly, a non-negative matrix factorization (NMF) framework was first constructed for spatial transcriptomic (ST) data (GEO: GSE225857; scCRLM: http://www.cancerdiversity.asia) based on a colorectal cancer tissue scRNA-seq reference dataset (GEO: GSE178341). A conditional auto-regressive (CAR) model was then applied to incorporate spatial constraints, enabling the estimation of posterior distributions for cell type abundance at each spot. Spots where CARD-inferred abundance of a specific phenotypic cell population exceeded the average level across all analyzed spots in the ST slice were designated cell spots for that phenotype. Subsequently, spatial co-localization patterns were assessed by calculating pairwise *Pearson* correlation coefficients between cell types, utilizing deconvolution results generated by the CARD algorithm as input.

##### Cell type co-occurrence analysis of single-cell resolution spatial transcriptomic data

Cellular composition and spatial organization within murine colon cancer tissues were analyzed using SeekSpace^®^ spatial transcriptomic sequencing technology at single-cell resolution. Following annotation of cellular phenotypes based on gene expression profiles, cells were spatially mapped, and co-occurrence analysis was subsequently performed using the Squidpy package. First, an adjacency graph was constructed from spatial coordinates (2D cellular positions), where nodes represented individual cells and edges denoted spatial neighborhood relationships. Co-occurrence scores for target subpopulations relative to other cell clusters were computed using the *sq.gr.co_occurrence* function. The co-occurrence score was defined as: co-occurrence score = p(exp cluster)/p(exp), quantifying the probability of observing neighboring cells from other subpopulations within the spatial vicinity of a specific cluster relative to their expected frequency across the entire tissue. Scores >1 indicated significant spatial co-occurrence (higher-than-random frequency), scores =1 reflected neutral spatial association (distribution matching global tissue patterns), and scores <1 suggested mutual exclusion or avoidance between subpopulations.

### QUANTIFICATION AND STATISTICAL ANALYSIS

#### Statistical and quantitative analysis of staining and microscopy images

For the statistical analysis of the confocal fluorescence images, we first quantified the relevant information using ImageJ software and then performed graphical and statistical analysis using PRISM. Differences were tested by using a one-way ANOVA followed by a Tukey’s multiple comparisons test (\**p* < 0.05; \*\**p* < 0.01, \*\*\**p* < 0.001).

#### Other statistical analysis

The data statistic was performed using PRISM. Paired or unpaired Student’s *t*-test (two tails) was used to compare experiments with only two groups. One-way ANOVA with multiple comparisons was performed for experiments with more than two groups. Kaplan-Meier assay and Log-rank test were used for survival analysis (ns: no significance; \**p* < 0.05; \*\**p* < 0.01, \*\*\**p* < 0.001).

## Notes

### Competing Interest Statement

The authors have declared no competing interest.

## References

2 Liu, Y. et al. Immune phenotypic linkage between colorectal cancer and liver metastasis. Cancer cell 40, 424–437 e425, doi:10.1016/j.ccell.2022.02.013 (2022).

3 Caronni, N. et al. IL-1beta(+) macrophages fuel pathogenic inflammation in pancreatic cancer. Nature 623, 415–422, doi:10.1038/s41586-023-06685-2 (2023).

4 Zhao, T., Luo, Y., Sun, Y. & Wei, Z. Characterizing macrophage diversity in colorectal malignancies through single-cell genomics. Front Immunol 16, 1526668, doi:10.3389/fimmu.2025.1526668 (2025).

5 Molgora, M. et al. TREM2 Modulation Remodels the Tumor Myeloid Landscape Enhancing Anti-PD-1 Immunotherapy. Cell 182, 886–900 e817, doi:10.1016/j.cell.2020.07.013 (2020).

6 Bill, R. et al. CXCL9:SPP1 macrophage polarity identifies a network of cellular programs that control human cancers. Science 381, 515–524, doi:10.1126/science.ade2292 (2023).

7 Mo, C. K. et al. Tumour evolution and microenvironment interactions in 2D and 3D space. Nature 634, 1178–1186, doi:10.1038/s41586-024-08087-4 (2024).

8 Ning, J. et al. Macrophage-coated tumor cluster aggravates hepatoma invasion and immunotherapy resistance via generating local immune deprivation. Cell Rep Med 5, 101505, doi:10.1016/j.xcrm.2024.101505 (2024).

9 Tan, J. X. & Finkel, T. A phosphoinositide signalling pathway mediates rapid lysosomal repair. Nature 609, 815–821, doi:10.1038/s41586-022-05164-4 (2022).

10 Rong, S. et al. DGAT2 inhibition blocks SREBP-1 cleavage and improves hepatic steatosis by increasing phosphatidylethanolamine in the ER. Cell metabolism 36, 617–629 e617, doi:10.1016/j.cmet.2024.01.011 (2024).

11 Lv, Q. et al. CSF1R inhibition reprograms tumor-associated macrophages to potentiate anti-PD-1 therapy efficacy against colorectal cancer. Pharmacol Res 202, 107126, doi:10.1016/j.phrs.2024.107126 (2024).

12 Ma, Y. & Zhou, X. Spatially informed cell-type deconvolution for spatial transcriptomics. Nat Biotechnol 40, 1349–1359, doi:10.1038/s41587-022-01273-7 (2022).

13 Feng, Y. et al. Spatially organized tumor-stroma boundary determines the efficacy of immunotherapy in colorectal cancer patients. Nature communications 15, 10259, doi:10.1038/s41467-024-54710-3 (2024).

14 La Manno, G. et al. RNA velocity of single cells. Nature 560, 494–498, doi:10.1038/s41586-018-0414-6 (2018).

15 Canellas-Socias, A., Sancho, E. & Batlle, E. Mechanisms of metastatic colorectal cancer. Nat Rev Gastroenterol Hepatol 21, 609–625, doi:10.1038/s41575-024-00934-z (2024).

16 Bala, P. et al. Aberrant cell state plasticity mediated by developmental reprogramming precedes colorectal cancer initiation. Science advances 9, eadf0927, doi:10.1126/sciadv.adf0927 (2023).

17 Li, T. et al. TIMER2.0 for analysis of tumor-infiltrating immune cells. Nucleic Acids Res 48, W509–W514, doi:10.1093/nar/gkaa407 (2020).

18 Li, H. et al. Dysfunctional CD8 T Cells Form a Proliferative, Dynamically Regulated Compartment within Human Melanoma. Cell 176, 775–789 e718, doi:10.1016/j.cell.2018.11.043 (2019).

19 Kamimoto, K. et al. Dissecting cell identity via network inference and in silico gene perturbation. Nature 614, 742–751, doi:10.1038/s41586-022-05688-9 (2023).

20 Lu, M. et al. Activation of the human chemokine receptor CX3CR1 regulated by cholesterol. Science advances 8, eabn8048, doi:10.1126/sciadv.abn8048 (2022).

21 Zhou, X. et al. Inhibition of DUSP18 impairs cholesterol biosynthesis and promotes anti-tumor immunity in colorectal cancer. Nature communications 15, 5851, doi:10.1038/s41467-024-50138-x (2024).

22 Su, P. et al. Enhanced Lipid Accumulation and Metabolism Are Required for the Differentiation and Activation of Tumor-Associated Macrophages. Cancer research 80, 1438–1450, doi:10.1158/0008-5472.CAN-19-2994 (2020).

23 Newman, A. M. et al. Robust enumeration of cell subsets from tissue expression profiles. Nat Methods 12, 453–457, doi:10.1038/nmeth.3337 (2015).

24 Sawada, N. et al. Transfer and Enzyme-Mediated Metabolism of Oxidized Phosphatidylcholine and Lysophosphatidylcholine between Low- and High-Density Lipoproteins. Antioxidants (Basel) 9, doi:10.3390/antiox9111045 (2020).

25 Zhang, F. et al. Inhibiting PLA2G7 reverses the immunosuppressive function of intratumoral macrophages and augments immunotherapy response in hepatocellular carcinoma. J Immunother Cancer 12, doi:10.1136/jitc-2023-008094 (2024).

26 Giurisato, E. et al. Extracellular-Regulated Protein Kinase 5-Mediated Control of p21 Expression Promotes Macrophage Proliferation Associated with Tumor Growth and Metastasis. Cancer research 80, 3319–3330, doi:10.1158/0008-5472.CAN-19-2416 (2020).

27 Tymoszuk, P. et al. In situ proliferation contributes to accumulation of tumor-associated macrophages in spontaneous mammary tumors. Eur J Immunol 44, 2247–2262, doi:10.1002/eji.201344304 (2014).

28 Liao, G. B. et al. Regulation of the master regulator FOXM1 in cancer. Cell Commun Signal 16, 57, doi:10.1186/s12964-018-0266-6 (2018).

29 Xu, R. et al. Invasive FoxM1 phosphorylated by PLK1 induces the polarization of tumor-associated macrophages to promote immune escape and metastasis, amplified by IFITM1. J Exp Clin Cancer Res 42, 302, doi:10.1186/s13046-023-02872-1 (2023).

30 Balli, D. et al. Foxm1 transcription factor is required for macrophage migration during lung inflammation and tumor formation. Oncogene 31, 3875–3888, doi:10.1038/onc.2011.549 (2012).

31 Yang, Y., Zhang, B., Yang, Y., Peng, B. & Ye, R. FOXM1 accelerates wound healing in diabetic foot ulcer by inducing M2 macrophage polarization through a mechanism involving SEMA3C/NRP2/Hedgehog signaling. Diabetes Res Clin Pract 184, 109121, doi:10.1016/j.diabres.2021.109121 (2022).

32 Liu, Y. H. et al. Functional macrophages and gastrointestinal disorders. World J Gastroenterol 24, 1181–1195, doi:10.3748/wjg.v24.i11.1181 (2018).

33 Casanova-Acebes, M. et al. Tissue-resident macrophages provide a pro-tumorigenic niche to early NSCLC cells. Nature 595, 578–584, doi:10.1038/s41586-021-03651-8 (2021).

34 Aziz, A., Soucie, E., Sarrazin, S. & Sieweke, M. H. MafB/c-Maf deficiency enables self-renewal of differentiated functional macrophages. Science 326, 867–871, doi:10.1126/science.1176056 (2009).

35 Liu, M. et al. Transcription factor c-Maf is a checkpoint that programs macrophages in lung cancer. The Journal of clinical investigation 130, 2081–2096, doi:10.1172/JCI131335 (2020).

36 Ecker, J. et al. The Colorectal Cancer Lipidome: Identification of a Robust Tumor-Specific Lipid Species Signature. Gastroenterology 161, 910–923 e919, doi:10.1053/j.gastro.2021.05.009 (2021).

37 Stuart, T. et al. Comprehensive Integration of Single-Cell Data. Cell 177, 1888–1902 e1821, doi:10.1016/j.cell.2019.05.031 (2019).

38 Bergen, V., Lange, M., Peidli, S., Wolf, F. A. & Theis, F. J. Generalizing RNA velocity to transient cell states through dynamical modeling. Nat Biotechnol 38, 1408–1414, doi:10.1038/s41587-020-0591-3 (2020).

39 Wolf, F. A., Angerer, P. & Theis, F. J. SCANPY: large-scale single-cell gene expression data analysis. Genome Biol 19, 15, doi:10.1186/s13059-017-1382-0 (2018).

40 Palla, G. et al. Squidpy: a scalable framework for spatial omics analysis. Nat Methods 19, 171–178, doi:10.1038/s41592-021-01358-2 (2022).

41 Menden, K. et al. Deep learning-based cell composition analysis from tissue expression profiles. Science advances 6, eaba2619, doi:10.1126/sciadv.aba2619 (2020).

42 Korsunsky, I. et al. Fast, sensitive and accurate integration of single-cell data with Harmony. Nat Methods 16, 1289–1296, doi:10.1038/s41592-019-0619-0 (2019).

43 Wu, T. et al. clusterProfiler 4.0: A universal enrichment tool for interpreting omics data. Innovation (Camb) 2, 100141, doi:10.1016/j.xinn.2021.100141 (2021).

44 Robinson, J. T. et al. Integrative genomics viewer. Nat Biotechnol 29, 24–26, doi:10.1038/nbt.1754 (2011).

45 Van Gassen, S. et al. FlowSOM: Using self-organizing maps for visualization and interpretation of cytometry data. Cytometry A 87, 636–645, doi:10.1002/cyto.a.22625 (2015).

46 Zi, R. et al. Metabolic-Immune Suppression Mediated by the SIRT1-CX3CL1 Axis Induces Functional Enhancement of Regulatory T Cells in Colorectal Carcinoma. Adv Sci (Weinh) 12, e2404734, doi:10.1002/advs.202404734 (2025).

47 Chen, S., Zhou, Y., Chen, Y. & Gu, J. fastp: an ultra-fast all-in-one FASTQ preprocessor. Bioinformatics 34, i884–i890, doi:10.1093/bioinformatics/bty560 (2018).

