## Supplementary data for "Metabolic Immunosuppression Mediated by Proliferative CD168^+^ TAMs in Colon Cancer"

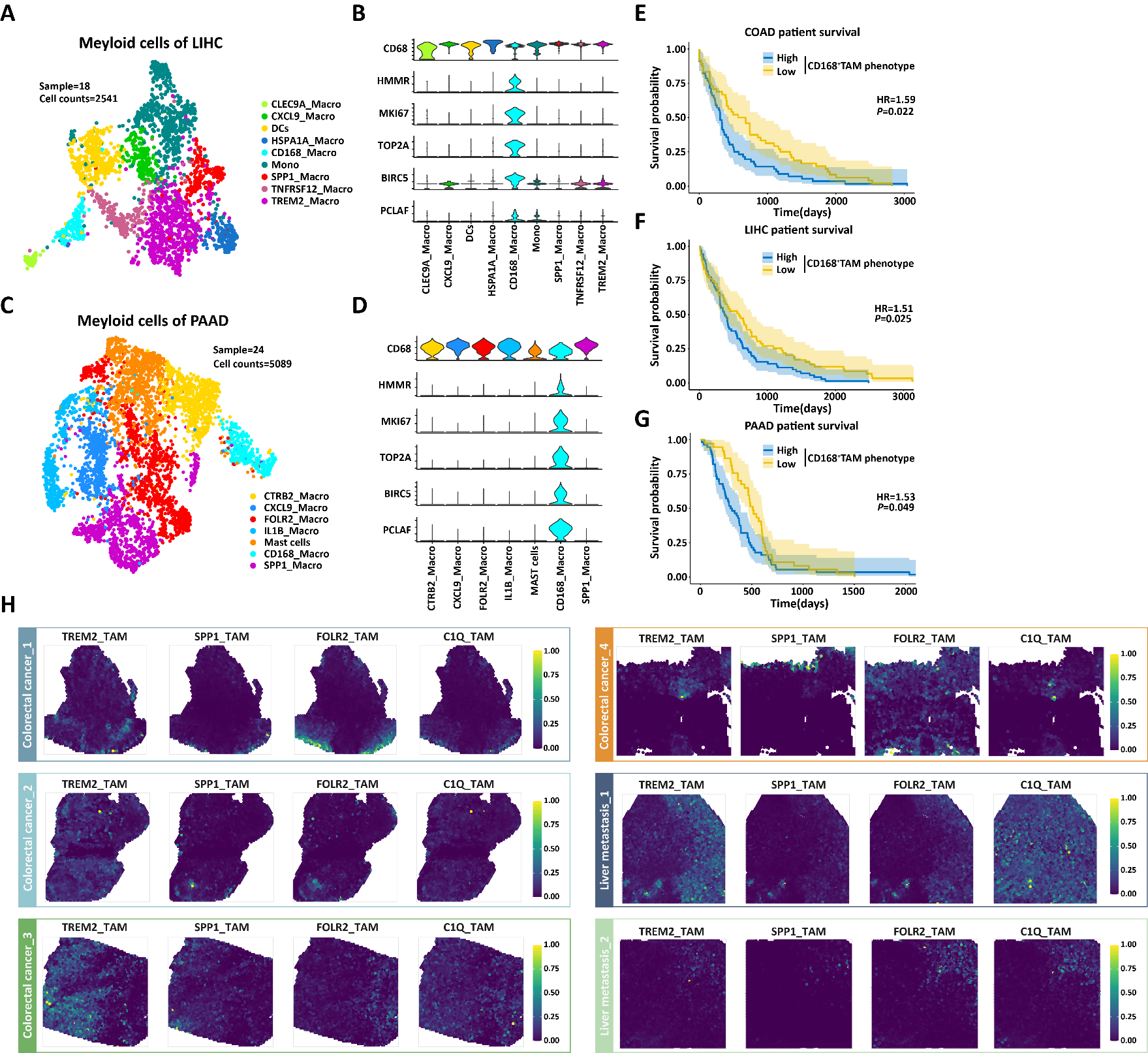
**Supplementary data Figure 1**

**Supplementary data Figure 1. Pan-cancer analysis of CD168^+^ TAM phenotypes and their association with clinical prognosis.**

(A) UMAP of scRNA-seq data (China National GeneBank Database: CNP0000650) of myeloid cells isolated from LIHC patients (n=18).

(B) Gene signatures of the CD168^+^ TAMs from LIHC.

(C) UMAP of scRNA-seq data (Genome Sequence Archive: CRA001160) of myeloid cells isolated from PAAD patients (n=24).

(D) Gene signatures of the CD168^+^ TAMs from PAAD.

(E-G) Kaplan–Meier survival analysis of patients with COAD, LIHC, and PAAD (from TCGA data) stratified by CD168^+^TAMs gene signature expression levels (normalized to CD68). Hazard ratio (HR) and P value of the Cox regression fit are shown.

(H) The spatial mapping of CARD-inferred illustrative cell types across samples.

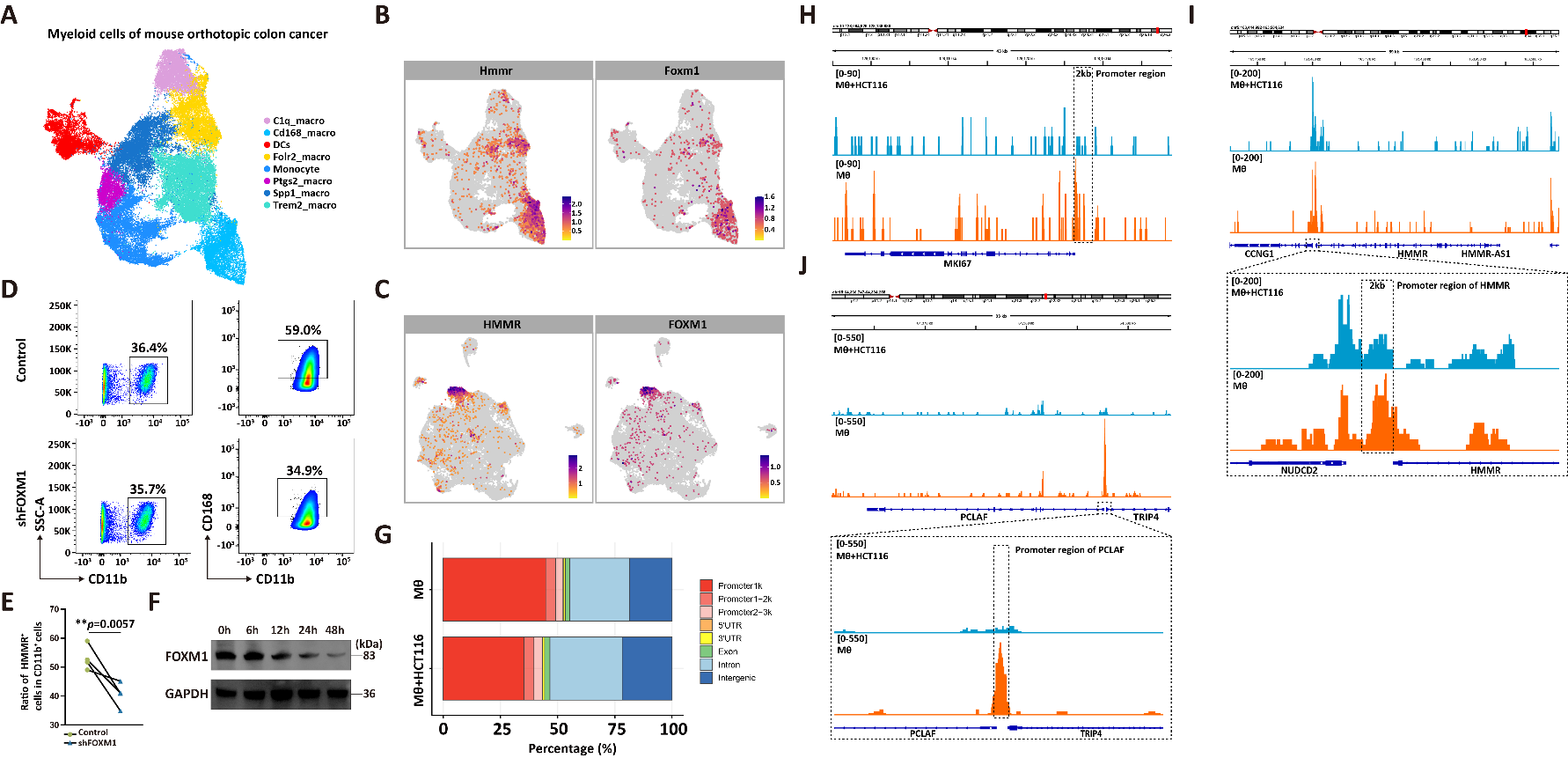
**Supplementary data Figure 2**

**Supplementary data Figure 2. FOXM1 is essential for sustaining the CD168^+^ TAMs phenotype.**

(A) UMAP of scRNA-seq data of Cd11b^+^ cells isolated from mouse orthotopic colon cancer.

(B) Comparative analysis of Hmmr and Foxm1 expression in Cd11b^+^ cells within murine colon cancer tissues.

(C) Comparative analysis of HMMR and FOXM1 expression in myeloid cells within human colon cancer tissues.

(D-E) Flow cytometric analysis of CD168^+^ phenotypic changes upon *FOXM1* knockdown in PBMC-derived and in vitro activated human macrophages, unpaired Student’s two-tailed t-test.

(F) Western blot analysis of FOXM1 expression levels in macrophages following co-culture with tumor cells in a Transwell system.

(G) Genomic occupancy of FOXM1 interaction sites in control macrophages versus HCT116-co-cultured macrophages.

(H-J) CUT&Tag sequencing analysis of FOXM1 enrichment at the *MKI67*, *HMMR*, and *PCLAF* promoter region in control macrophages versus HCT116-co-cultured macrophages.

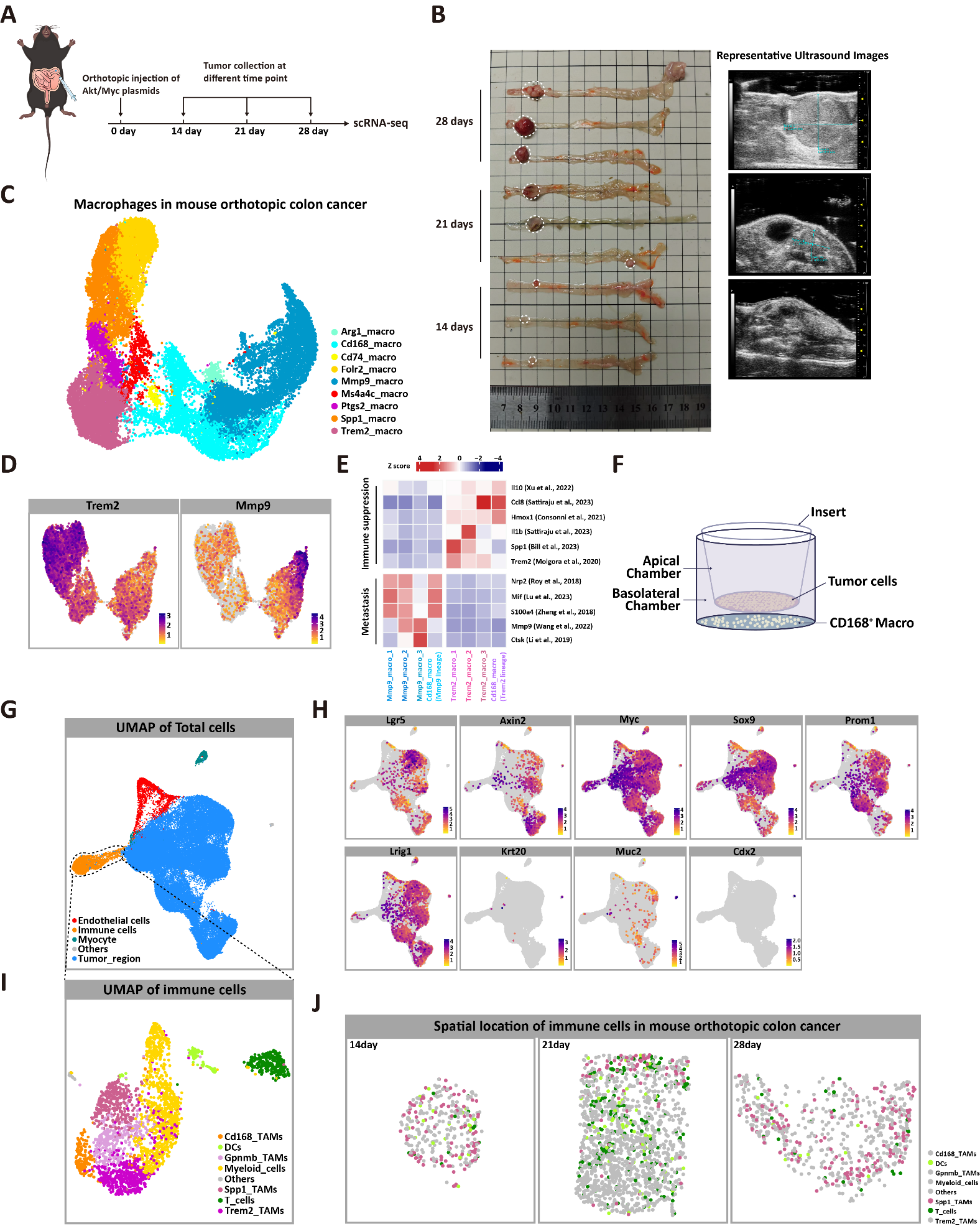
**Supplementary data Figure 3**

**Supplementary data Figure 3. Single-cell transcriptomic and single-cell spatial transcriptomic analyses of mouse orthotopic colon cancer.**

(A) Schematic diagram of the construction and sampling of the tumor tissues in the mouse orthotopic colon cancer model.

(B) Left: Murine colon tumor specimens from distinct time points. Right: Representative *in vivo* ultrasound images of corresponding groups (n = 3 mice/group).

(C) UMAP of scRNA-seq data of macrophages from mouse orthotopic colon cancer.

(D) Expression of Trem2 and Mmp9 in Cd168^+^ TAM-derived polarized cells.

(E) Differential expression profiles of immunosuppressive and pro-metastatic genes in Trem2 and Mmp9 lineage TAMs.

(F) Schematic representation of CD168^+^ macrophage-tumor cell co-culture in a Transwell system.

(G) UMAP of spatial transcriptomic data of cells from mouse orthotopic colon cancer.

(H) Expression of established Apc^KO^-induced colon cancer marker genes in primary tumor cells derived from our experimental model.

(I) UMAP of spatial transcriptomic data of immune cells from mouse orthotopic colon cancer.

(J) Spatial mapping of immune cells in the mouse orthotopic colon cancer (Highlight Spp1^+^ TAMs, T cells, and DC cells).

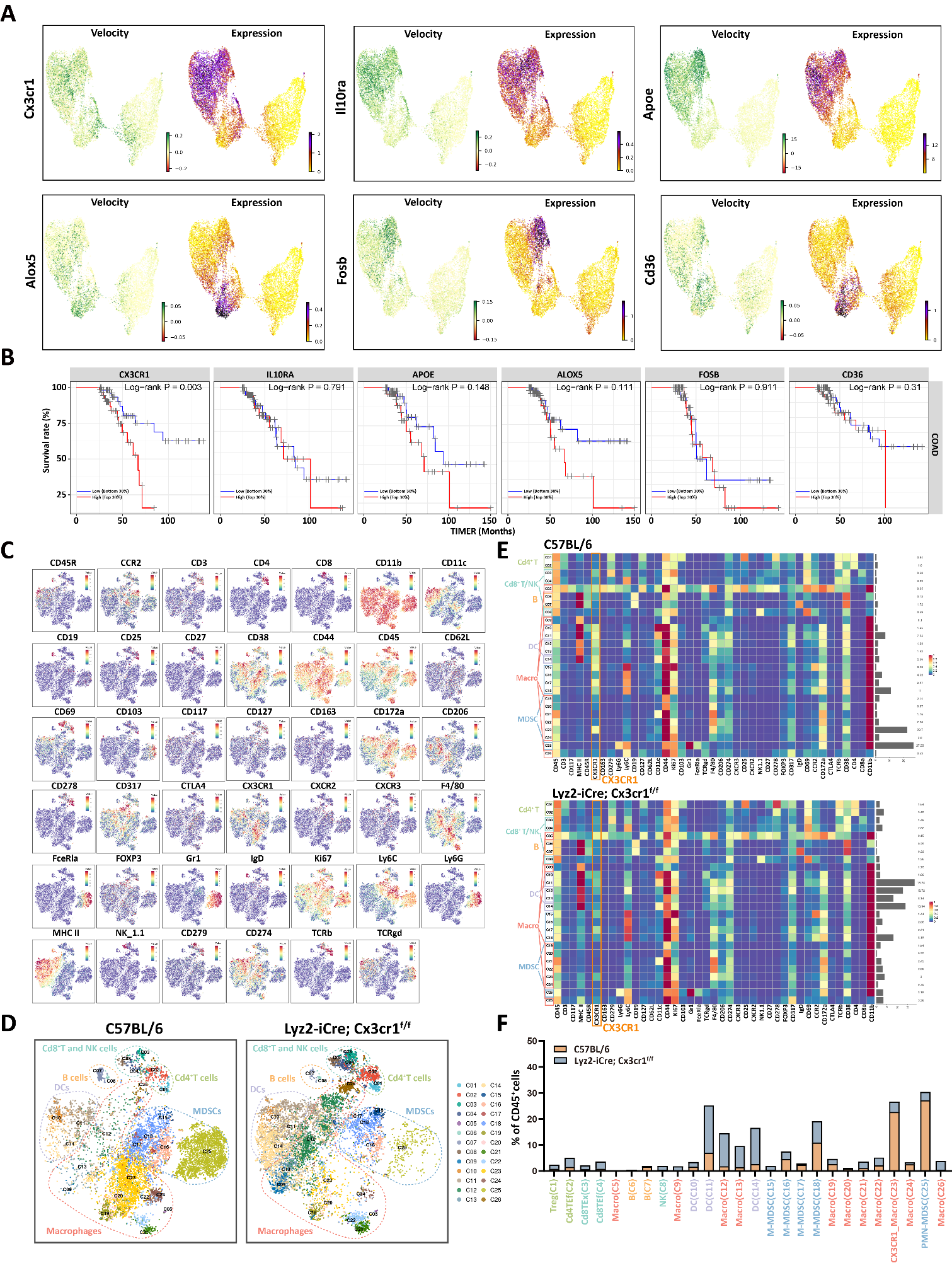
**Supplementary data Figure 4**

**Supplementary data Figure 4. The CX3CR1 signaling promotes the immunosuppressive polarization of TAMs.**

(A) Expression dynamics and expression level of putative driver genes underlying Trem2 lineage polarization of Cd168^+^ TAMs.

(B) Kaplan-Meier survival analysis of colon cancer samples from the TCGA cohort, stratified by median expression levels of *Cx3cr1*, *Il10ra*, *Apoe*, *Alox5*, *Fosb*, and *Cd36*.

(C) CD45^+^ immune cells were isolated from orthotopic colon tumors of C57BL/6 (n=3) and Lyz2-iCre; Cx3cr1^f/f^ (n=3) mice, pooled separately, and subjected to CyTOF analysis. The synchronized expression patterns of 41 functionally annotated protein markers across all single cells were displayed.

(D) t-SNE projection of 6 distinct immune subsets from C57BL/6 and Lyz2-iCre; Cx3cr1^f/f^ mice.

(E) Heatmap shows the expression of the 41 markers in different cell populations, with CX3CR1 highlighted to show its conditional knockout in the myeloid cells of Lyz2-iCre; Cx3cr1^f/f^ mice.

(F) Proportions of immune cell subsets among total CD45^+^ cells were quantified across experiment groups.

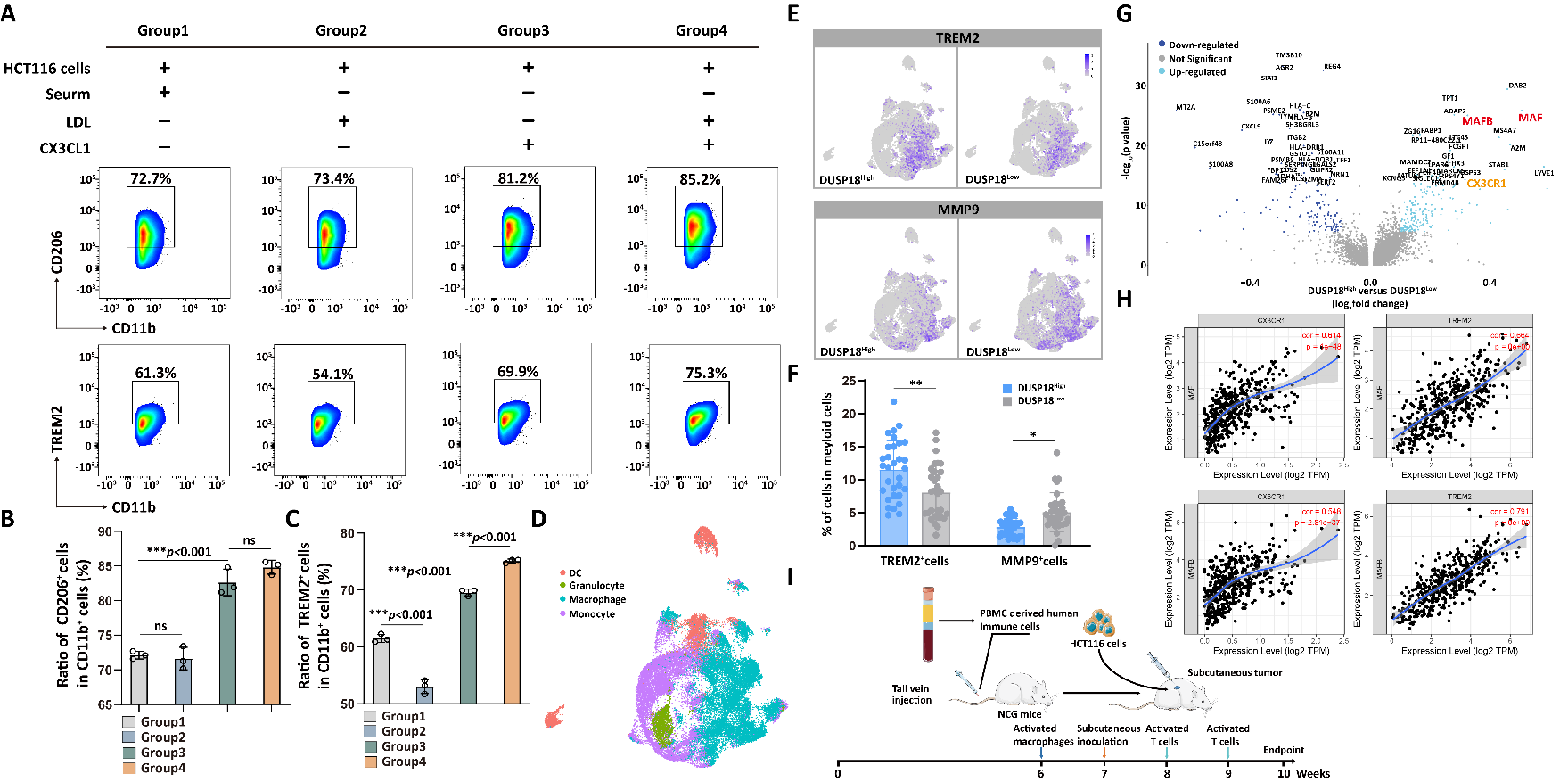
**Supplementary data Figure 5**

**Supplementary data Figure 5. Cholesterol molecules synergistically activate CX3CR1 with CX3CL1.**

(A-C) CD168^+^ macrophages were co-cultured with HCT116 cells in a Transwell system for 48 hours following the illustrated treatment. Contour plots and frequencies of TREM2^+^ (CD11b^+^TREM2^+^) and M2-type (CD11b^+^CD206^+^) TAMs in different groups, one-way ANOVA.

(D) UMAP visualization of myeloid cell subpopulations from scRNA-seq data of CRC tissues (GEO: GSE178341).

(E) Comparative analysis of TREM2 and MMP9 expression in macrophages from DUSP18^High^ versus DUSP18^Low^ cohorts.

(F) Frequencies of TREM2^+^ and MMP9^+^ macrophages in DUSP18^High^ and DUSP18^Low^ samples, unpaired Student’s two-tailed t-test.

(G) Volcano plot depicting DEGs in CD168^+^ TAMs between DUSP18^High^ and DUSP18^Low^ samples.

(H) Spearman correlation analysis was performed to evaluate associations between depicted gene expression profiles in colon adenocarcinoma (COAD) samples from the TCGA database.

(I) Schematic showing the construction of the humanized subcutaneous xenograft mouse model.

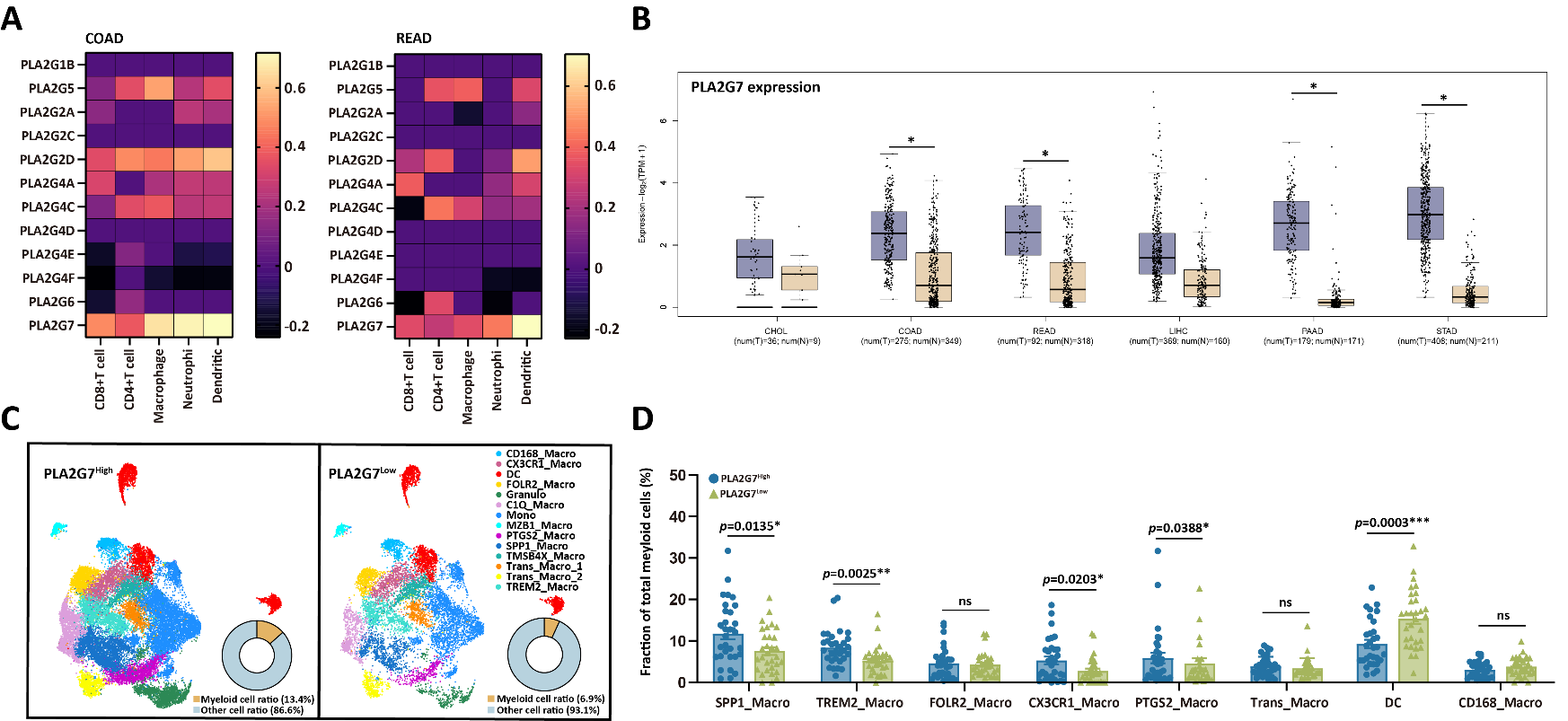
**Supplementary data Figure 6**

**Supplementary data Figure 6. PLA2G7 expression modulates myeloid infiltration and phenotypic dynamics in CRC.**

(A) Immune cell infiltration in TCGA colorectal cancer samples was quantified using the CIBERSORT algorithm. Associations between immune cell abundance and expression levels of key metabolic enzymes in the glycerophospholipid pathway were analyzed via Pearson correlation.

(B) Comparative PLA2G7 mRNA expression in primary digestive tract malignancies and paired adjacent normal tissues (TCGA RNA-seq data, Wilcoxon rank-sum test, **p* < 0.001).

(C) UMAP plot distinct myeloid cell clusters in scRNA-seq data of CRC tissues (GEO: GSE178341) stratified by PLA2G7 expression levels; the accompanying pie chart quantifies myeloid cell prevalence within the total cellular population.

(D) The differential composition of myeloid cell subpopulations between PLA2G7-High and PLA2G7-Low expression cohorts from the CRC scRNA-seq data, Wilcoxon rank-sum test.

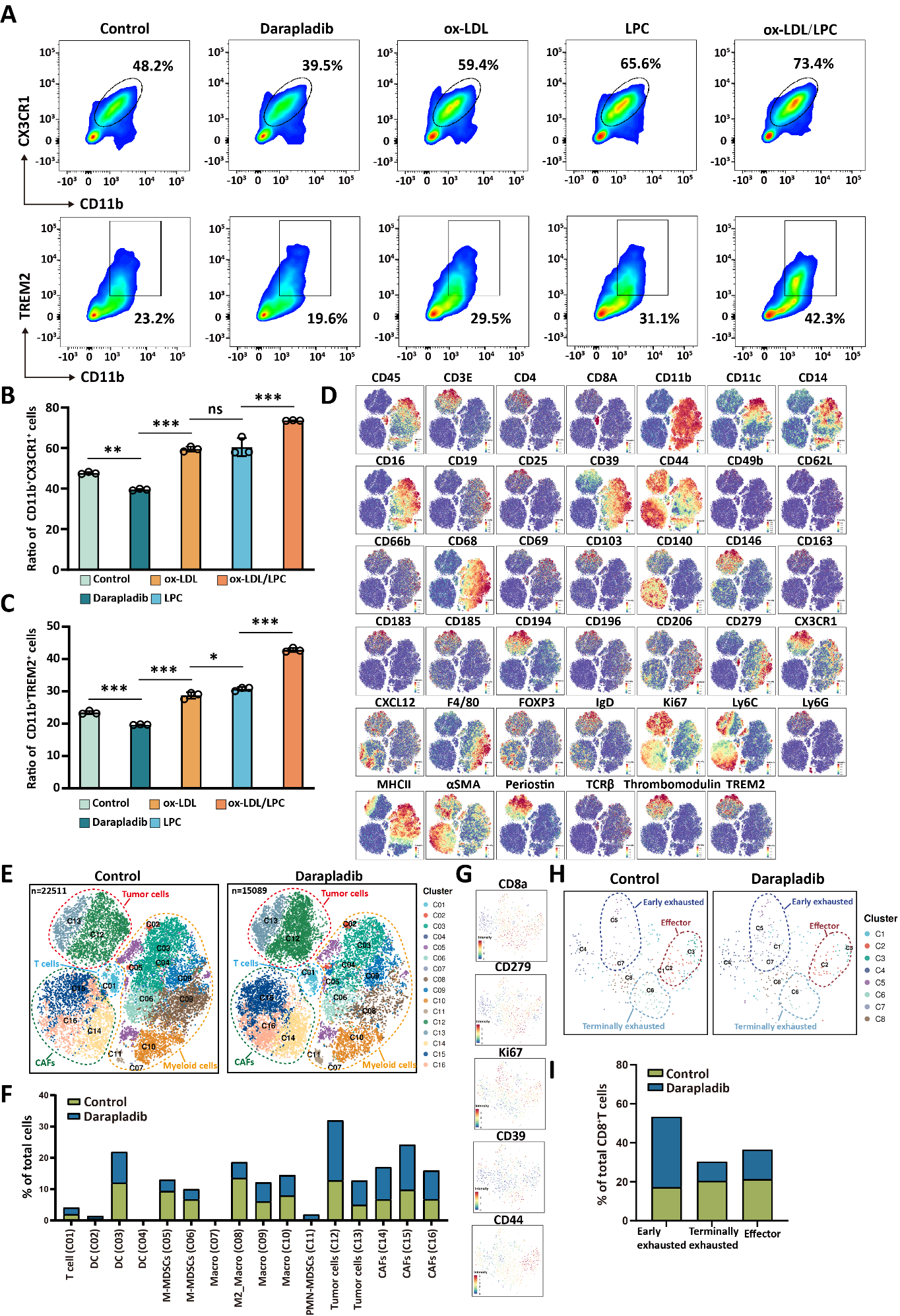
**Supplementary data Figure 7**

**Supplementary data Figure 7. Inhibition of PLA2G7 reprograms the tumor immune microenvironment in colon cancer.**

(A-C) PBMC-derived macrophages and HCT116 cells were co-cultured in a Transwell system for 48 hours under the illustrated conditions. Contour plots and frequencies of CX3CR1^+^ (CD11b^+^CX3CR1^+^) and TREM2^+^ (CD11b^+^TREM2^+^) TAMs across experimental groups, one-way ANOVA.

(D) Mice received intraperitoneal administration of Darapladib (10 mg/kg) every other day, commencing 7 days post-intracolonic oncogene injection. Treatment continued until the experimental endpoint on day 21, when animals were euthanized for tumor tissue harvest. Orthotopic colon tumors of control (n=3) and Darapladib-treated (n=3) mice were collected, pooled separately, and subjected to CyTOF analysis. The synchronized expression patterns of 41 functionally annotated protein markers across all single cells were displayed.

(E) t-SNE projection of tumor, T cell, CAF, and myeloid cell subsets from control and Darapladib-treated mice.

(F) Proportions of cell subsets among total cells were quantified between the control and Darapladib-treated groups.

(G-H) CD8^+^ T cells were classified into three distinct subsets based on marker expression profiles: effector cells (CD8^+^CD44^+^PD-1^-^CD39^-^Ki67^+^), early-stage exhausted cells (CD8^+^CD44^−^PD-1^+^CD39^+^Ki67^+^), and late-stage exhausted cells (CD8^+^CD44^−^PD-1^+^CD39^+^Ki67^−^).

(I) Proportions of cell subsets among total CD8^+^ T cells were quantified between the control and Darapladib-treated groups.

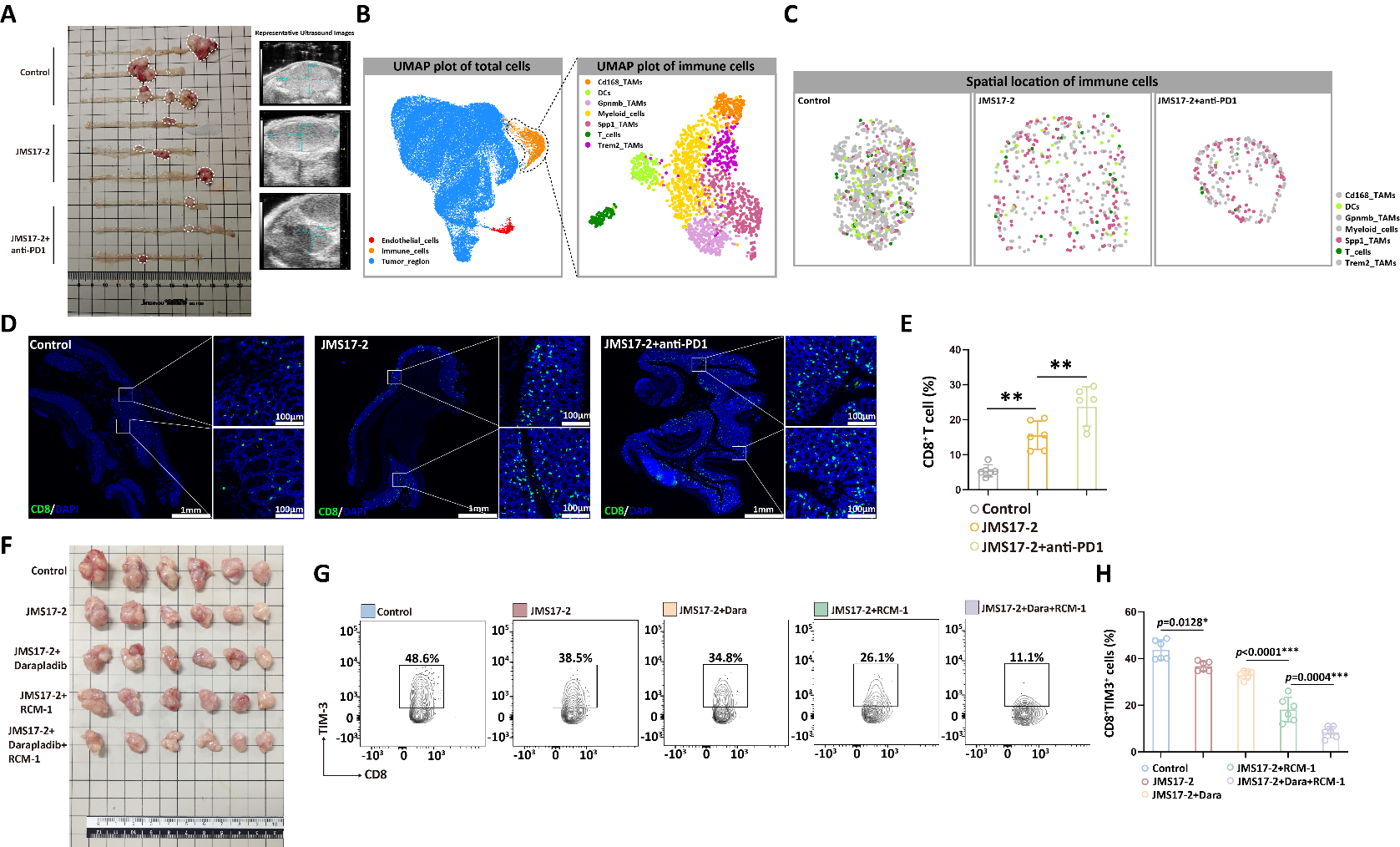
**Supplementary data Figure 8**

**Supplementary data Figure 8. CD168^+^ TAMs targeting reprogram the immune landscape in colon cancer.**

(A) Left: Murine colon tumor specimens from distinct experimental cohorts. Right: Representative *in vivo* ultrasound images of corresponding treatment groups (n = 3 mice/group).

(B) Mice were orthotopically injected with oncogene plasmids to initiate tumorigenesis. Starting 7 days post-injection, experimental groups received either JMS17-2 (10 mg/kg daily) or anti-PD-1 (250 μg per mouse every 3 days) treatments for 14 consecutive days. All animals were sacrificed 21 days following plasmids administration for tumor collection and subsequent spatial transcriptomic analysis. UMAP of spatial transcriptomic data of total cells (left) and immune cells (right) from murine colon cancer.

(C) Spatial mapping of immune cells in the mouse orthotopic colon cancer across experimental groups (Highlight Spp1^+^ TAMs, T cells, and DC cells).

(D) Immunofluorescence analysis of CD8^+^ T cell infiltration in peritumoral tissues across experimental groups. Representative images show spatial distribution patterns of CD8^+^ T cells (green), scale bar=1mm (left); scale bar=100μm (right magnified view).

(E) Total cells (DAPI, blue) and CD8^+^ T cells (green) in tumor tissues were quantified using ImageJ to calculate the proportion of CD8^+^ T cells relative to total cellularity across experimental groups. Each experimental group comprised three samples. Two representative fields of view from mIF images were analyzed for each sample. Data represent mean ± SEM; **P < 0.01 by one-way ANOVA.

(F) Representative photographs of the tumors from humanized subcutaneous xenograft mice across experimental groups.

(G-H) Contour plots and frequencies of TIM-3^+^CD8^+^ T cells across experimental groups, one-way ANOVA.
